# Biocompatibility assessment of SilkMA-Gelatin ink tuned for 3D extrusion bioprinting

**DOI:** 10.1101/2025.11.09.687334

**Authors:** Prachi Agarwal, Jagnoor Singh Sandhu, Himanshu Kumar Bhatt, Paramita Das, Raviraja N Seetharam, Kirthanashri S Vasanthan

## Abstract

3D bioprinting applications have led to the development of multifunctional scaffolds that are biocompatible, overcoming shear stress to provide structural integrity. We report the isolation of silk fibroin (SF) using calcium chloride (CaCl_2_), which was further developed into an ink comprising silkMA and gelatin, demonstrating effective cytocompatibility and biocompatibility when 3D printed using an extrusion-based 3D bioprinter. Rheological characterization of SilkMA-Gelatin showed an improved viscosity and printability of the formulation as compared to an unmodified silk-gelatin composition. SilkMA-gelatin was bioprinted with NIH3T3 cells and demonstrated enhanced viability and proliferation, highlighting its cytocompatibility and ability to support a favourable 3D microenvironment. *In vivo* biocompatibility of the 3D bioprinted SilkMA-Gelatin was evaluated in Wistar rats, where 3D printed scaffolds maintained structural stability for up to 1 month, with no inflammatory response, progressive cell infiltration, and re-epithelialization. Serum biochemistry analysis, including ALP, ALT, LDH, and Urea (BUN), remained within normal physiological ranges, further confirming systemic safety. Based on the findings, the optimized SilkMA-gelatin ink proves to be highly effective in fabricating cell-laden structures that are stable, exhibit enhanced cytocompatibility and biocompatibility, and offer promising potential for advanced tissue regeneration applications.

**Graphical abstract:** Schematic overview of 3D bioprinting in wound healing and biomedical applications

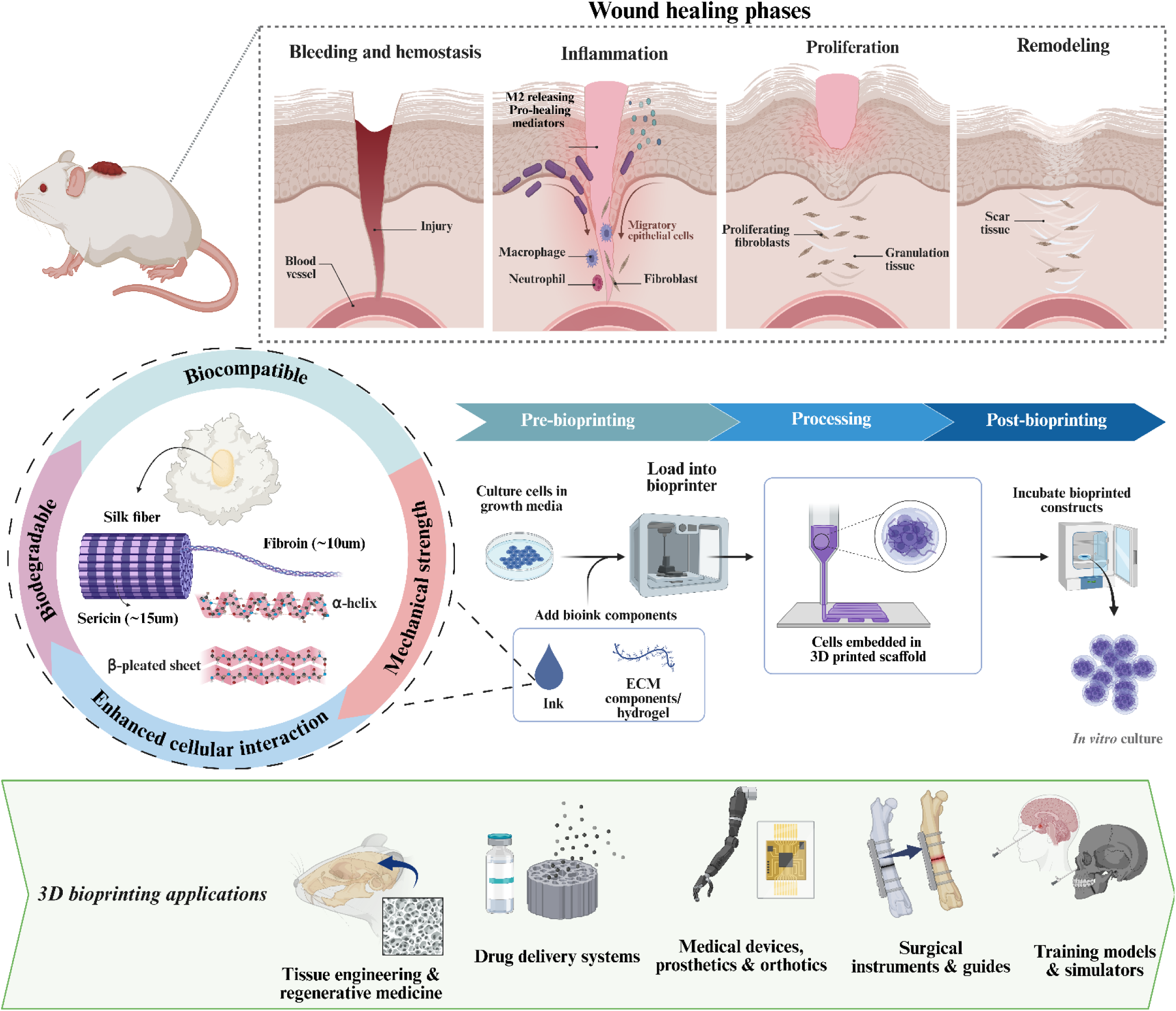

## 1. Introduction

Wound healing is a complex and well-orchestrated process of cellular and molecular events, involving sequential phases of hemostasis, inflammation, proliferation and tissue remodelling. The process starts as soon as there is an infliction of wound and the remodelling of the scar can take upto 1-2 years ^1^. Human body possesses a remarkable intrinsic ability to heal wounds, however in cases of chronic or severe wounds such as those resulting from diabetes, burns or infections the healing cascade becomes impaired ^2^.

When the natural repair mechanisms fail, wounds may remain open for prolonged periods thereby leading to risk of infection, necrosis, and in severe cases amputation^3^. One of the reasons of delayed healing can be the prolonged inflammation in case of chronic or burn wounds or underlying conditions of the body such as diabetes^4^, ulcers^5^, malnutrition^6^ etc. Absence of a bioactive, cell interacting matrix at the wound site further impairs the cellular migration and tissue regeneration which demands the need for innovative therapeutic effects^7^.

Three dimensional (3D) bioprinting is an advanced approach that integrates engineering, materials science, and biology to produce complex 3D objects. This technology allows fabrication of cell laden constructs that can mimic the native surrounding of the tissues. The technology also allows precise placement of the bioink in a layer-by-layer manner ensuring uniform distribution of cells within the polymer matrix, enabling the fabrication of 3D printed scaffolds that are customized to meet the anatomical and functional requirements of the patient^8^. Extrusion-based 3D bioprinting technique is known for its ability to handle various bioinks and supports the incorporation of multiple bioactive components including growth factors, extracellular matrix proteins or cells, creating spatially heterogenous constructs that mimics the gradients observed in the native tissues ^9^.

The ideal scaffolds utilized in tissue engineering should have excellent biocompatibility, biodegradability, mechanical integrity and enhanced porosity (>90%) and interconnectivity for better cell proliferation and adhesion^10^. Abiding by it, silk and gelatin, the two natural proteinaceous polymers possess the properties of being an ideal biomaterial. Other properties such as degradation time can be controlled or altered based on the specific features of tissue development^11^.

Silk fibroin derived primarily from domestic silkworm cocoons, such as *Bombyx mori*, is FDA-approved for biomedical use and composed of approximately 75-83.3% fibrous fibroin and 16.7-25% amorphous sericin^12^. It exhibits a hierarchical structure comprising of β-sheet and α-helices contributing to its exceptional mechanical strength, elasticity and toughness^13^. SF’s biocompatibility, minimal immunogenicity and tunable biodegradability make it an attractive material for the application in 3D printing. Presence of integrin binding motifs like RGD peptide sequences and hydrophobic domains that facilitate cell adhesion and ECM interaction^14^.

Hybrid biomaterials can be used to customize biodegradability to the application. While structurally similar to collagen, SF is commonly employed as a reinforcing component in hydrogels to increase mechanical strength^15^. Blending SF with a protein with inherent RGD sequences, such as gelatin, can increase silk hydrogel bioactivity. Gelatin is a soluble protein made from collagen, the major ECM component of connective tissues, including bone, cartilage, and skin, by partial acid (type A) ^16^. Silk-gelatin composite hydrogels are often fabricated by immobilizing gelatin within crosslinked silk networks using chemical or enzymatical component^17^.

Elaborating on this foundation, SilkMA-Gelatin 3D printed scaffold has been developed to harness the combined effectiveness of both the polymers, resulting in a 3D printed scaffold with enhanced printability, rheological behavior, excellent swelling and hydrophilic characteristics. Both *in vitro* and *in vivo* evaluation has been observed to establish cytocompatibility, biocompatibility and the potential of the silk-based 3D printed scaffold in cutaneous tissue regeneration. The novelty of this work is to lies in the integration of silkMA with gelatin and MTGase achieving a mechanically robust yet biologically responsive 3D printed scaffolds which can further be used for regenerative applications.

## 2. Materials and Methods

### 2.1 Materials

Silk cocoons were obtained from *Bombyx mori* purchased from the silk market, Bengaluru, India. Gelatin (porcine source), Type A, 175G, Phosphate Buffer Saline (PBS), and Glycidyl methacrylate (GMA) were purchased from Sigma Aldrich, India. Culture media Dulbecco’s Modified Eagle Medium (DMEM), Low glucose and DMEM-High glucose, and Fetal Bovine Serum (FBS) were procured from Hi-media, India. Antibiotics (Penicillin-streptomycin) and trypsin-EDTA used for cell culture were purchased from Gibco, India. Wistar rats were procured and housed in the Central Animal Research Facility (CARF), Manipal Animal Research Facility, Manipal, India. Live–dead cell viability kits were purchased from Invitrogen, USA. Biochemical assays such as Alkaline phosphatase (ALP) and Alanine transaminase (ALT) were purchased from Abcam. Lactate dehydrogenase (LDH) and Urea Nitrogen (BUN) were obtained from Elabsciences.

### 2.2 Animal Ethics Approval

All the experimental procedures involving animals were approved by the Institutional Animal Ethics Committee (IAEC), Manipal Academy of Higher Education, Manipal, as per CPSCEA guidelines (IAEC Registration No. 94/PO/RReBi/S/99/CPCSEA). Following ethical standards and guidelines set forth by the Animal Research: Reporting of *In Vivo* Experiments (ARRIVE) guidelines, rigorous protocols were adhered to in all studies involving laboratory animals.

### 2.3 Isolation of Silk Fibroin

#### 2.3.1 Degumming of silk

About 10 g of sliced cocoons were boiled in 300 mL of 0.5 M Na_2_CO_3_ for 1 hour at 100°C. Subsequently, the obtained degummed SF was washed thrice with distilled water, left in a fume hood overnight for 24 hours to dry at room temperature, and weighed to measure the total amount of sericin loss. The degumming loss associated with each degumming process is calculated using the equation^18^:

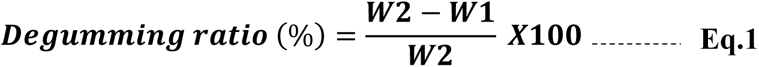

where W_1_ = weight of sample before degumming, W_2_ = weight of sample after degumming

#### 2.3.2 Extraction of SF & SilkMA Synthesis

About 20% (w/v) solution of degummed SF was prepared by immersing tightly packed degummed SF fibers in 5.5 M calcium chloride (CaCl_2_) solution. The beaker was sealed with aluminium foil and placed on a hot plate stirrer at a temperature of 90°C and stirred at 1000 rpm. Post the dissolution of fibers glycidyl methacrylate (GMA) was added at a rate of 0.5 mL/min to a final concentration of 10% (v/v). The mixture was then stirred at 1000 rpm for 6 hours at 60 °C. Following this, the solution was dialyzed against distilled water at 37°C for 3 days, using a cellulosic dialysis membrane (molecular weight cutoff 10-12 kDa, Thermo Scientific, USA)^19^. Finally, the obtained SilkMA solutions were lyophilized for 2 days, resulting in a white, porous foam stored at −80 °C until needed.

##### 2.3.2.1 Preparation of SilkMA-Gelatin Ink

An equal weight of lyophilized SilkMA and gelatin (type A, 175G) was stirred at 45 °C for 15 minutes at 330 rpm. About 500 µL/mL of MTGase (50U) was added to the ink and stirred for 15 minutes (330 rpm at 50°C). The obtained ink was further incubated for 15 minutes (50°C) before being stored at 4°C until use.

##### 2.3.2.2 Analytical characterization of SilkMA

The isolated SF and crosslinked SF (SilkMA) were subjected to FTIR to confirm the peak shift changes using a Bruker Alpha II. The spectral range was kept between 4000–550 cm^−120^.

##### 2.3.2.3 Thermal Characterization of SilkMA-Gelatin

Differential Scanning Calorimetry (DSC) was performed using a DSC (PerkinElmer DSC 6000) calorimeter. The samples were heated in sealed aluminium pans under nitrogen flow at a scanning rate of 5°C/min from 0.1 to 100°C/min^21^. The heat flow as a function of temperature was measured for SilkMA, gelatin, and SilkMA-Gelatin.

### 2.4 3D scaffold construction using CAD modelling system

The design of the 3D scaffolds was carried out using computer-aided design (CAD) software. Scaffold geometries were created using Autodesk Fusion, which allowed for parametric control over dimensions and porosity features. The final CAD models were exported in. stl format and subsequently sliced using the slicing software Cura to generate the G-code required for 3D bioprinting. The slicing parameters were optimized to ensure high print fidelity, uniform pore size, and consistent layer deposition. The G-code was then used to fabricate the scaffolds using the Anga Pro bioprinter, which operates on a pneumatic extrusion mechanism^22^. Layer-by-layer deposition paths were carefully optimized for structural integrity and reproducibility across batches. The final scaffold design incorporated layer-by-layer deposition paths optimized for print fidelity and structural integrity.

### 2.5 Material Characterization

#### 2.5.1 Rheological Characterization of Silk-based Ink

The storage modulus (*G*′) and loss modulus (*G*″) of Silk-gelatin and SilkMA-Gelatin inks were measured in the oscillatory mode at 37 °C using a plate–plate geometry and a 1.2 mm gap between plates (MCR 702 Multi Drive, Anton-Paar, Austria). After running an amplitude sweep to determine the linear viscoelastic range (LVER), a frequency sweep was conducted between 0.01 and 10 Hz at 0.5% shear strain^23^.

#### 2.5.2 Morphological characterization

The surface morphology of the 3D printed SilkMA-Gelatin samples was analysed using field emission scanning electron microscopy (FESEM) (JEOL make JSM-7610FPlus) (1.5kV accelerating voltage, 5mm working distance, 2 × 10^−4^ Pa vacuum pressure)^24^.

#### 2.5.3 Swelling Properties

The dry weight of SilkMA-Gelatin 3D printed SilkMA-Gelatin scaffolds (1cm^3^) was noted. They were soaked in PBS (pH 7.4) at a 37 °C incubator. At predetermined time points from 1 hour to 4 days, the scaffolds were taken out, and excess PBS was removed, and the swelling ratio (SR) was calculated^25^:

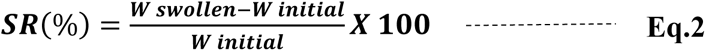

#### 2.5.4 Wettability analysis

Surface wettability was measured using a contact angle meter (Holmarc’s Contact Angle Meter; Model No: HO-IAD-CAM 01). The sessile drop method was used to estimate the water contact angles (CA) of the 3D printed scaffolds^26^.

### 2.6 Cytocompatibility assays

SilkMA-Gelatin ink was prepared as mentioned in section [2.3.2.3] and subsequently loaded with NIH3T3 cells (1x10^5^ cells per 3D bioprinted scaffold) using a sterile connector and syringe under minimal pressure. Post-cell incorporation, the bioink was 3D bioprinted into a scaffold of 1cm^3^ using Anga Pro. The 3D bioprinted constructs were incubated in high glucose DMEM at 37°C with 5% CO_2_ and cultured at 1,7, and 14 days to evaluate cytocompatibility. The 3D bioprinted scaffolds were refreshed with culture media every two days.

#### 2.6.1 Live/Dead Staining

The cytocompatibility of the 3D bioprinted scaffolds was assessed using the Live/Dead assay kit according to the manufacturer’s instructions. At predetermined time points of day 1, 7, and 14, the 3D bioprinted scaffold were transferred to another well, washed with PBS, and stained using the live/dead cell imaging kit. Post incubation, scaffolds were observed using an Evos M7000 fluorescent microscope (Thermo Fisher Scientific, USA) for live (green) and dead (red) cells^27^.

#### 2.6.2 Hemotoxylin & Eosin staining (H&E staining)

The 3D bioprinted scaffolds at day 1, 7, and 14 were fixed in 10% formaldehyde and further dehydrated for paraffin embedding. Cryosectioning was performed, and sections of the 3D bioprinted scaffolds were sliced. The initial few sections were discarded, and approximately at 50 slices, the sections were collected^28^. H&E staining was performed for the sections of 3D bioprinted scaffolds to check the infiltration of NIH3T3 cells in the scaffolds. The sections were further visualised using an Evos M7000 microscope (Thermo Fisher Scientific, USA).

### 2.7 Biocompatibility assays

#### 2.7.1 Implantation

Male Wistar rats (9–12-week-old) weighing 150-200g were obtained for the biocompatible *in vivo* studies. The animals were randomly divided into three groups (Control, Sham, SilkMA-Gelatin) before surgery. Rats were anesthetized via an intraperitoneal injection of 10mg/kg xylazine and 90mg/kg ketamine. The skin was shaved and disinfected with 70% alcohol. A full-thickness wound measuring approximately 1.5 cm x 1.5 cm x 1.5 cm was created on the dorsum of the animals using a sterile surgical blade^29^. SilkMA-Gelatin scaffolds were implanted in the pouch created in the skin. The scaffold consisted of five layers, with a total height of 1.05 mm. Each layer has a thickness of 0.075 mm, ensuring uniform stacking and structural integrity. The sutured area was covered with a standard dressing protocol. The animals were sacrificed at 14 and 30 days to evaluate the impact of the implanted 3D scaffold.

#### 2.7.2 Hematological parameters evaluation

Blood sample was collected from the Wistar rats before surgery at day 14 and day 30. The samples were collected via a retro-orbital sinus and then immediately transferred to 0.5mL EDTA-coated tubes, and Complete Blood Count (CBC) was measured using a Celltac hematology analyzer (Nihon Kohden, Japan)^30^. The mean ± SD of hematological parameters, including White Blood Cells (WBCs), Lymphocytes, Red Blood Cells (RBCs), Platelets, and Hematocrit (HCT) was evaluated according to the normal physiological range referenced from^31^.

#### 2.7.3 Biochemical assays

The collected blood is immediately transferred into a serum collection tube (without anticoagulant) and allowed to clot at room temperature for 30–60 minutes. After clot formation, the sample is centrifuged at 2000g for 10–15 minutes to separate the serum from the cellular components. The clear supernatant (serum) is carefully pipetted into sterile microcentrifuge tubes and stored at -20°C or -80°C for long-term preservation^32^. Proper handling and storage are crucial to prevent hemolysis, which can interfere with biochemical analyses such as enzyme activity, hormone levels, and metabolic markers.

To evaluate the chronic biological response to the 3D printed scaffolds, biochemical assays including blood urea (BUN), alanine aminotransferase (ALT), alkaline phosphatase (ALP), and lactate dehydrogenase (LDH) were performed^32^.

#### 2.7.4 H&E staining

After animal sacrifice (14 & 30 days), tissue samples were excised from the area of the wound and liver were collected and immediately fixed in 4% paraformaldehyde^33^. The obtained samples were then embedded in paraffin for sectioning using a cryostat (Leica CM 1860UV), and slices ranging from 8 to 16μm thick were cryosectioned and stained with H&E.

### 2.8 Statistical analysis

All the experiments were performed in triplicate, and the results were reported as mean ± standard deviation. The data analysis was performed using OriginPro, Microsoft Excel, and GraphPad Prism 9, with analysis of variance (ANOVA). P-values less than 0.05 were considered statistically significant.

## 3. Results and Discussion

### 3.1 Physicochemical Characterization

Silk cocoons were degummed using 0.5 M Na_2_CO_3_ and NaHCO_3_ by boiling them for 1 hour at 90°C, and washed thoroughly with distilled water **(Figure 1)**. A higher degumming efficiency was observed with Na_2_CO_3_ (27.39 ± 0.349 %) compared to NaHCO_3_ (23.9 ± 1.5 %), as shown in **(Figure 2a)**. SEM micrographs **(Figure 2b)** highlight the morphological changes in silk cocoons. Silk cocoons exhibited a layered, fibrillar surface with a smoother and more uniform fibre structure showing minimal disruption in case of Na_2_CO_3_. In contrast, NaHCO_3_-treated cocoons displayed numerous filaments that were tightly packed with sericin. Na_2_CO_3_ being a stronger base compared to NaHCO_3,_ is a better degumming agent as it creates a more alkaline environment, enhancing rapid breaking of hydrogen bonds and peptide interactions in sericin leading to its hydrolysis followed by removal ^34^.

**Figure 1:**
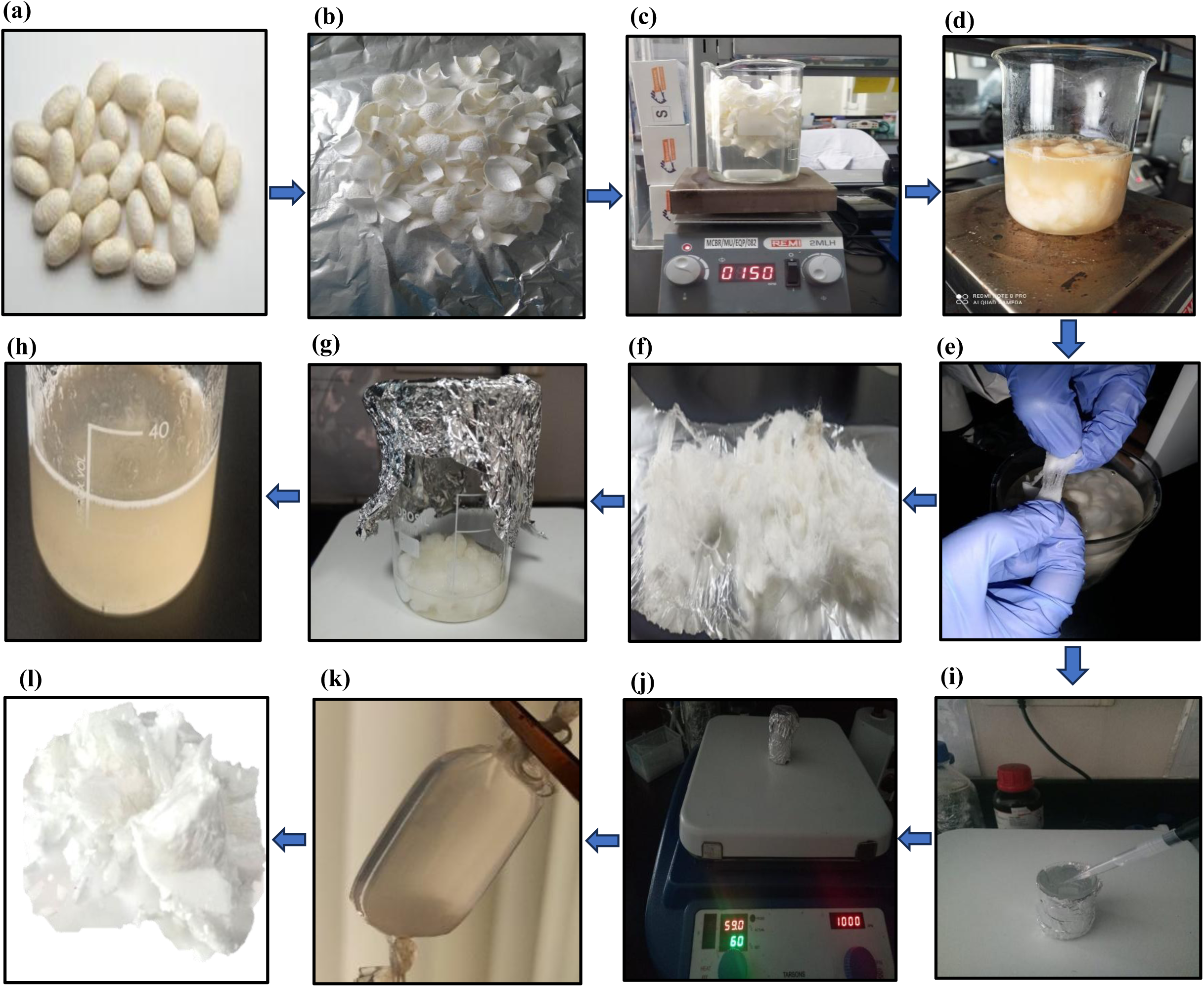
(a) Silk cocoons obtained from *Bombyx mori,*(b) Cutting of silk cocoons in small pieces for effective degumming, (c) Silk cocoons in Na_2_CO_3_ solution (0 hour), (d) Silk cocoons in Na_2_CO_3_ solution (0.5 hour), (e) Silk cocoons in Na_2_CO_3_ solution ( 1 hour), (f) Degummed silk, (g) Silk fibroin (SF) isolation in calcium chloride (CaCl_2_) solution (0 hour), (h) Isolated silk fibroin (3 hours), (i) Addition of GMA for crosslinking, (j) Crosslinking for 6 hours at 1000rm at 60°C, (k) Dialysis of SilkMA, (l) Lyophilized SF

**Figure 2:**
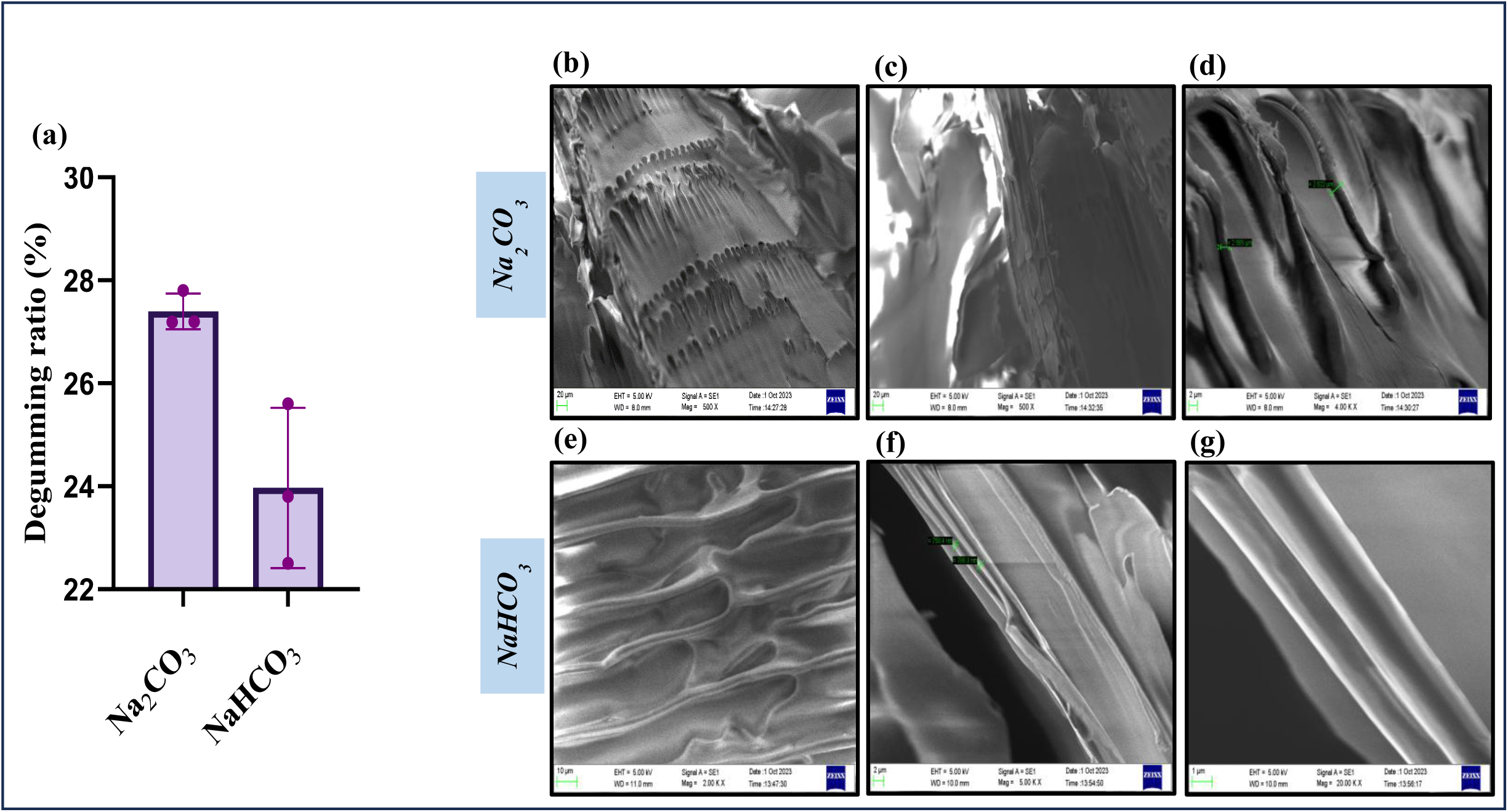
(a) Comparison of degumming efficiency using Na₂CO₃ and NaHCO₃, The degumming ratio (%) of silk fibers treated with Na₂CO₃ and NaHCO₃, (b) Scanning Electron Micrograph acquired at 5 kV at 500X magnification, (c) 2000X, (d) 4000X, (e) 2000 X, (f) 5000X, (g) 20,000 X representing the silk fibers after degumming

### 3.2 Preparation of SilkMA-Gelatin

During the preparation of ink as depicted in **Figure 3**, SF was initially crosslinked using GMA. It was observed at that at various concentrations and despite varying the gelation hours, stable gels were not obtained. To ensure a stable structure with enhanced integrity, type A gelatin (175 G) and MTGase was added to obtain stable ink, which will enhance the functionality and facilitate 3D printing. **Table 1** presents various combinations of SilkMA-Gelatin to obtain stable printability with shear stress behaviour.

**Figure 3:**
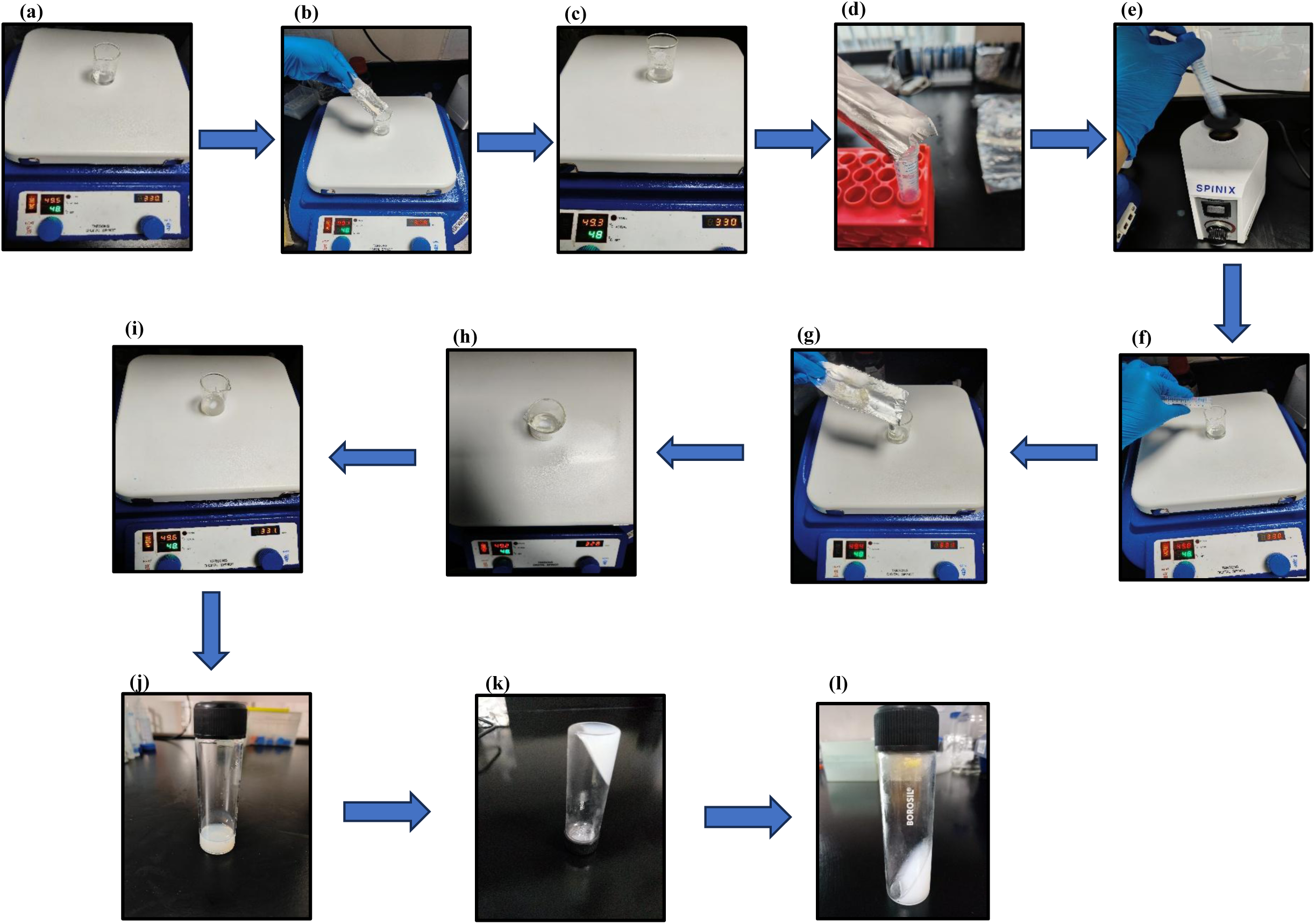
Schematic representation of SilkMA-Gelatin ink preparation: PBS was maintained at 48°C and 330 rpm. (a-c) 7% (w/v) gelatin (Type A, 175G) was dissolved until transparent, (e-f) Preparation of MTGase solution, vortexed and added to the gelatin solution, followed by stirring, (g-i) Addition of SilkMA & stirred until dissolved, (j) SilkMA-Gelatin solution incubated at 50°C, (k-l) Gelation of SilkMA-Gelatin ink

**Table 1:**
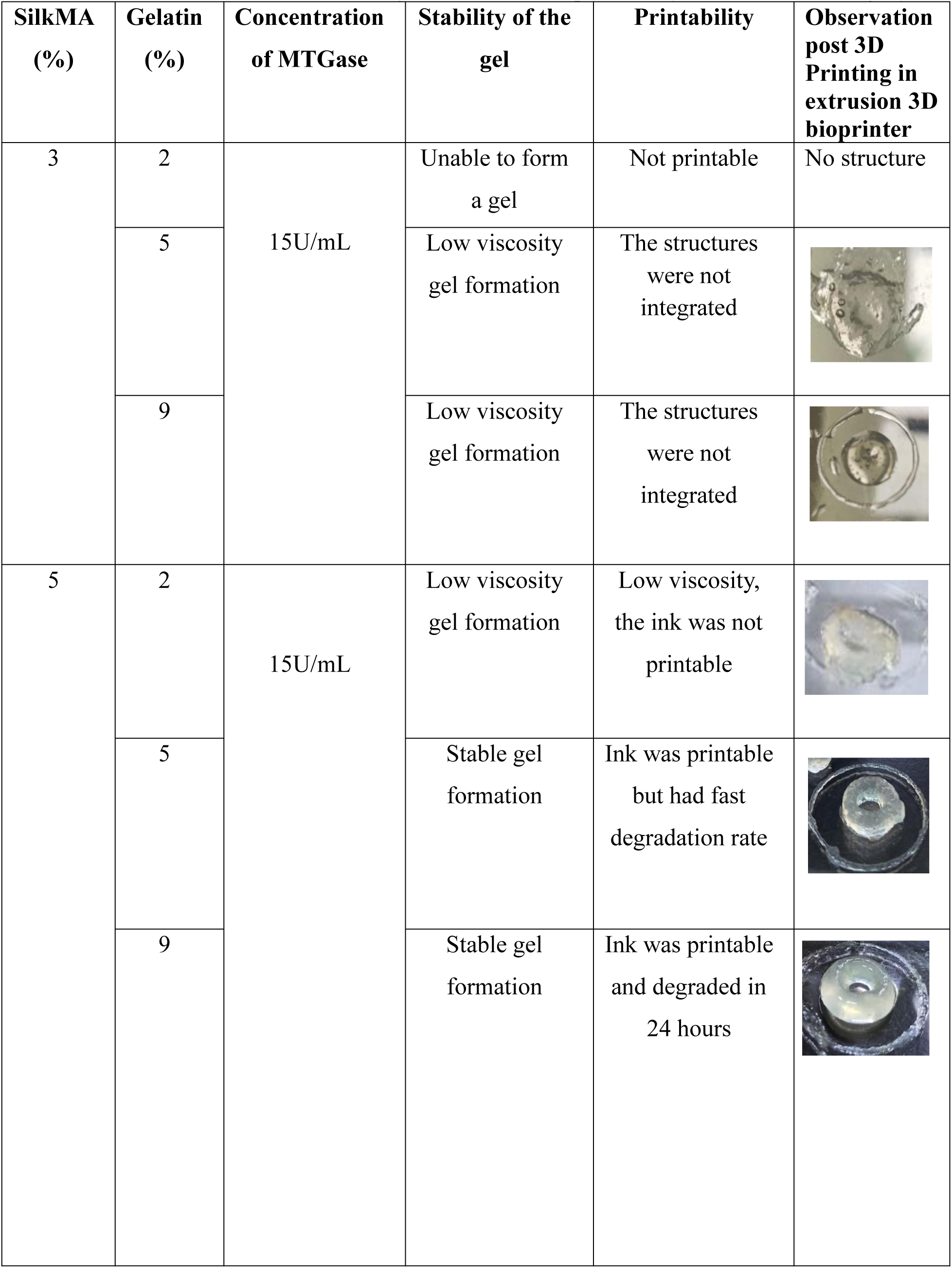
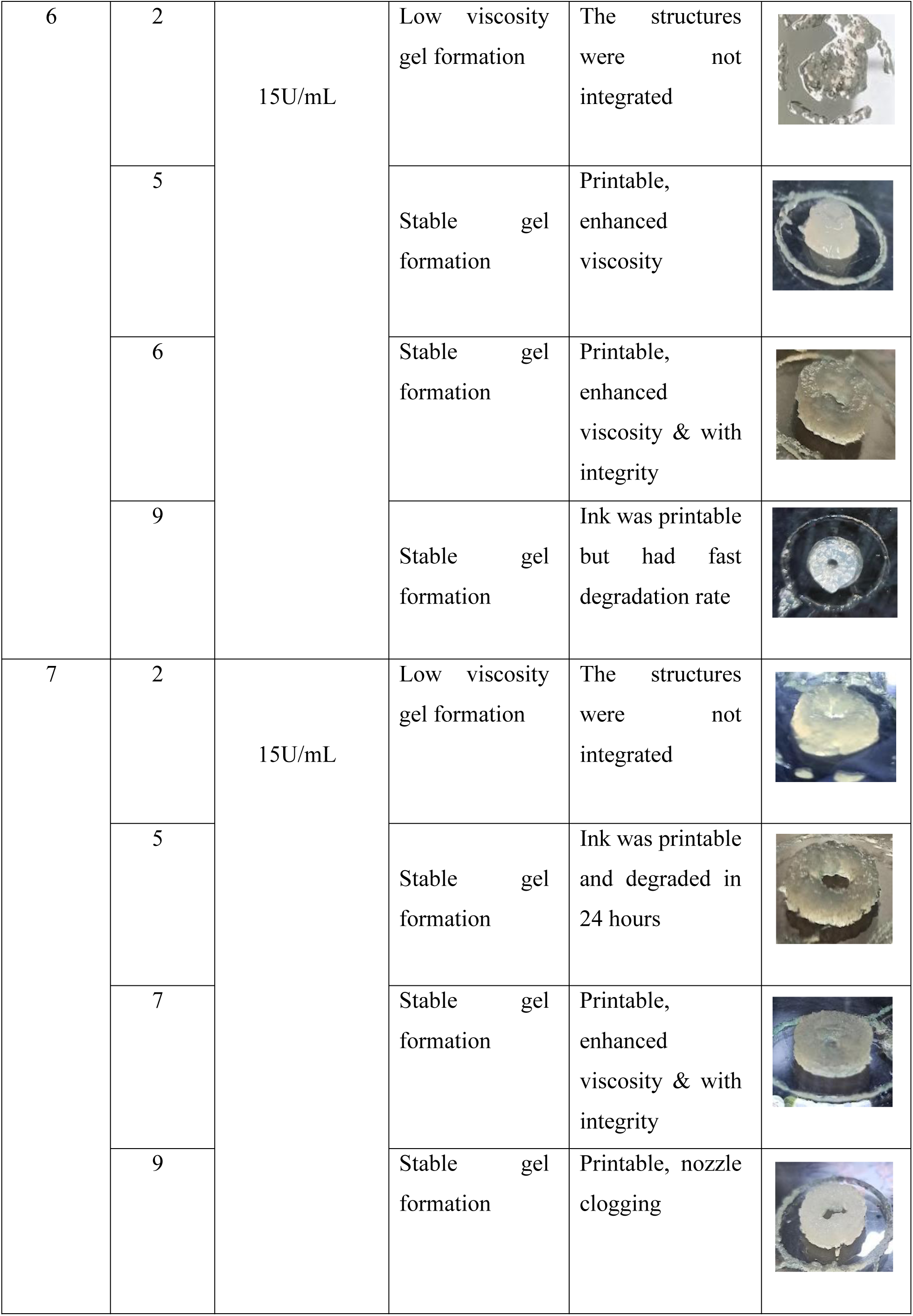
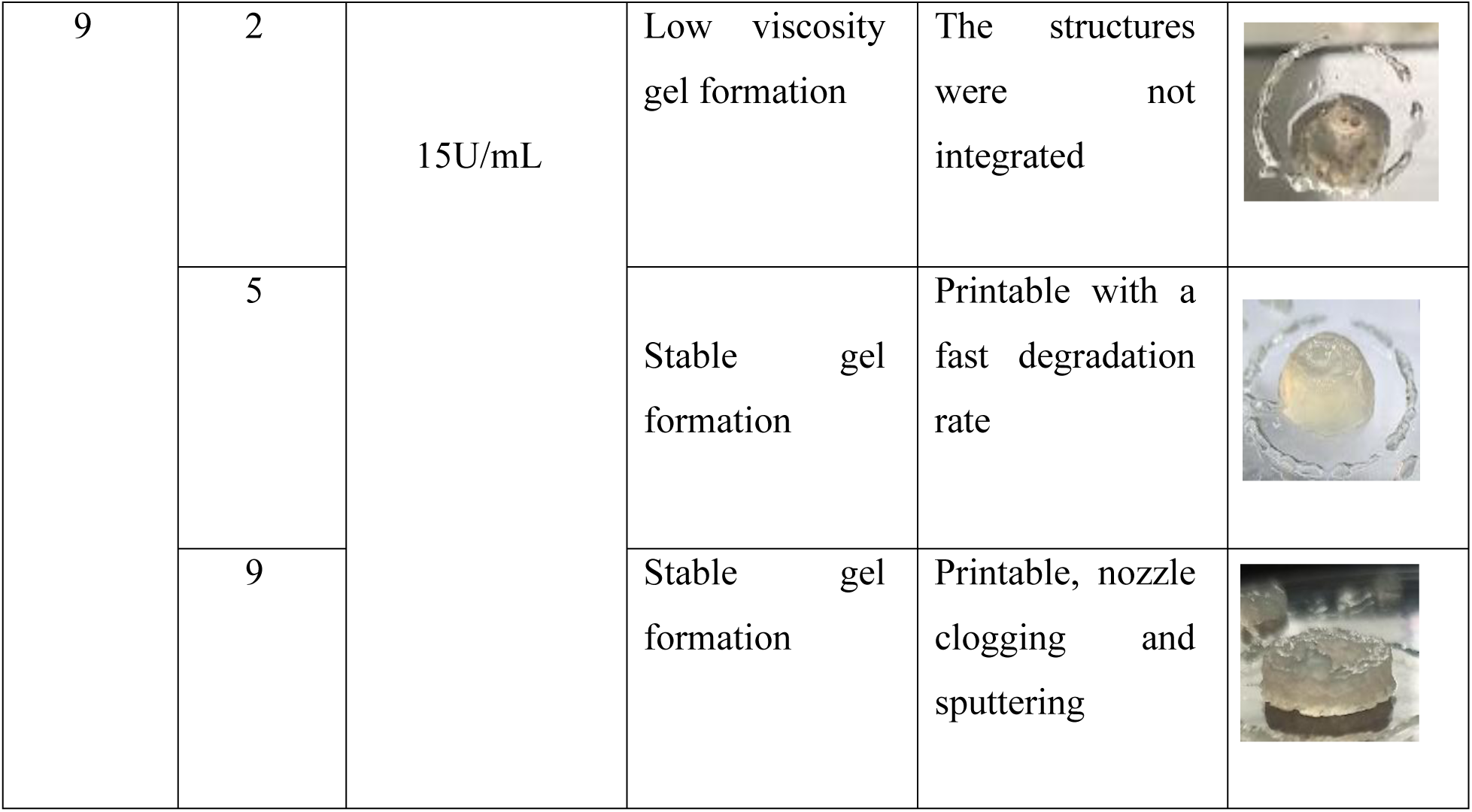
Optimization of SilkMA-Gelatin formulations with varying gelatin concentrations with constant SilkMA and MTGase ratios. Representative images of the 3D printed constructs are illustrated to demonstrate effect of gelatin content on printability and structural integrity

| **SilkMA (%)** | **Gelatin (%)** | **Concentration of MTGase** | **Stability of the gel** | **Printability** | **Observation post 3D Printing in extrusion 3D bioprinter** |
| --- | --- | --- | --- | --- | --- |
| 3 | 2 | 15U/mL | Unable to form a gel | Not printable | No structure |
|  | 5 |  | Low viscosity gel formation | The structures were not integrated |  |
|  | 9 |  | Low viscosity gel formation | The structures were not integrated |  |
| 5 | 2 | 15U/mL | Low viscosity gel formation | Low viscosity, the ink was not printable |  |
|  | 5 |  | Stable gel formation | Ink was printable but had fast degradation rate |  |
|  | 9 |  | Stable gel formation | Ink was printable and degraded in 24 hours |  |
| 6 | 2 | 15U/mL | Low viscosity gel formation | The structures were not integrated |  |
|  | 5 |  | Stable gel formation | Printable, enhanced viscosity |  |
|  | 6 |  | Stable gel formation | Printable, enhanced viscosity & with integrity |  |
|  | 9 |  | Stable gel formation | Ink was printable but had fast degradation rate |  |
| 7 | 2 | 15U/mL | Low viscosity gel formation | The structures were not integrated |  |
|  | 5 |  | Stable gel formation | Ink was printable and degraded in 24 hours |  |
|  | 7 |  | Stable gel formation | Printable, enhanced viscosity & with integrity |  |
|  | 9 |  | Stable gel formation | Printable, nozzle clogging |  |
| 9 | 2 | 15U/mL | Low viscosity gel formation | The structures were not integrated |  |
|  | 5 |  | Stable gel formation | Printable with a fast degradation rate |  |
|  | 9 |  | Stable gel formation | Printable, nozzle clogging and sputtering |  |

Optimization of gelatin and MTGase concentrations was carried out to achieve a formulation which has enhanced printability, structural integrity and stability for 3D printing and bioprinting applications. The physical characteristics of the ink were highly dependent on the relative ratios of silkMA and gelatin. At lower concentrations 2-3%, the formulation exhibited inadequate viscosity and poor layer fidelity resulting in non-printable structures. Similar behavior was observed for unbalanced ratios (5:2,7:2,9:2, 9:5) (5:2,7:2,9:2, 9:5) which failed to maintain the structural definition post-deposition due to insufficient polymer-polymer interactions necessary for network stabilization.

Intermediate concentration ranging from 5-7% demonstrated optimal physical properties with continuous extrusion and good shape retention, indicating an appropriate balance between viscosity and shear thinning behavior. At higher concentrations (8-9%) the viscosity increased excessively, leading to nozzle clogging and irregular extrusion. This aligns with the previous findings where excessive polymer content enhances crosslinking density but negatively affects flow behavior and printability ^35^.

For the initial optimization phase cylindrical scaffolds were printed due to simple geometry and uniform cross-section which facilitated evaluation of extrusion behavior, filament continuity and layer stacking. Cylindrical scaffolds serve as a practical geometrical structure to observe the rheological variations in filament uniformity, spreading or layer fusion the circular periphery allowing quick identification of formulations that lacks sufficient viscosity to create a 3D model ^36^.

For the wound healing applications, cubical scaffolds are more spatially compatible architecture in providing contact with planar tissue surfaces allowing uniform distribution of pressure and ensuring consistent interface adhesion to the wound bed^37^. From a mechanical standpoint, cubical scaffolds constructs provide a balanced stress profile during handling & dressing changes, thereby reducing deformation as compared to the cylindrical scaffolds^37^. Thus, cubical SilkMA-Gelatin scaffolds were 3D printed for further optimizations.

After the concentrations were further optimized for printing, degradation was the next challenge, which was further troubleshooted using an increased concentration of 50U/mL of MTGase demonstrated in **Table 2**, demonstrating that 7% of SilkMA-Gelatin ink has enhanced printability, structural integrity, and maintains a degradation rate of over 1 month. It was also observed that with 15U/mL of MTGase the degradation of the 3D printed scaffolds was within 12-24 hours probably due to insufficient crosslinking. With higher concentration of MTGase (50 U/mL) it was observed that the printability along with the structural integrity and degradation rate of the scaffolds improved significantly to 1 month and conclusively equal ratio of SilkMA and Gelatin (7%) with 50U/mL of MTGase provided the most desirable balance of physical, mechanical, rheological and functional properties. The reason behind an increased concentration of MTGase is that it leads to enhanced enzymatic crosslinking due to the catalysing effect of MTGase forming ε-(γ-glutamyl) lysine bonds between lysine and glutamine residues, resulting in a stable, three-dimensional network. This process helps in minimizing the gelation time, increasing stifness of the gel and reduced biodegradation due to tight molecular packing and decreased chain mobility^38^.

**Table 2:**
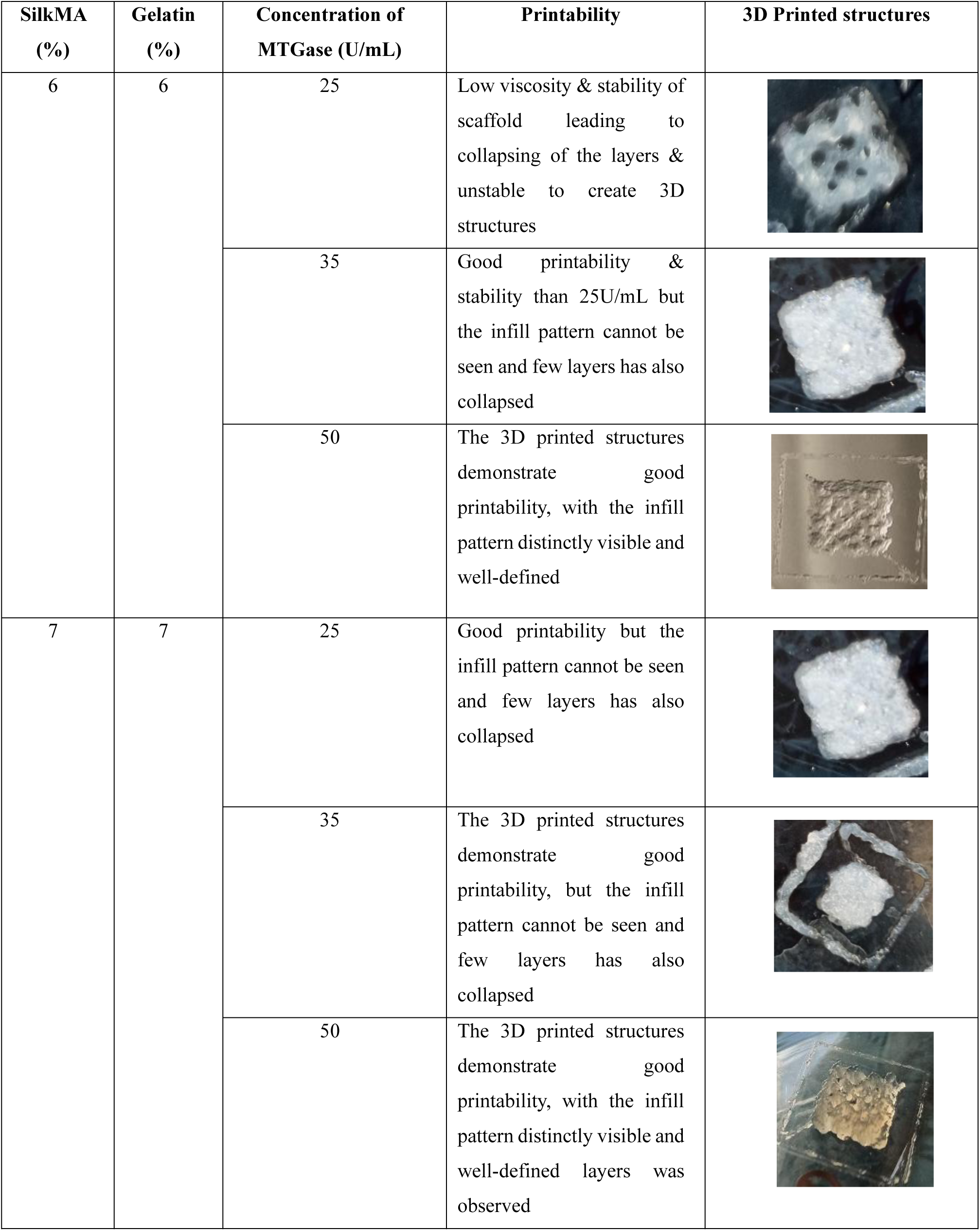
Optimization of SilkMA-Gelatin ink formulation prepared with equal concentration of SilkMA and Gelatin while varying the concentration of MTGase. Images illustrate the effect of MTGase crosslinking on printability and shape retention of 3D printed SilkMA-Gelatin scaffolds

| SilkMA (%) | Gelatin (%) | Concentration of MTGase (U/mL) | Printability | 3D Printed structures |
| --- | --- | --- | --- | --- |
| 6 | 6 | 25 | Low viscosity & stability of scaffold leading to collapsing of the layers & unstable to create 3D structures |  |
|  |  | 35 | Good printability & stability than 25U/mL but the infill pattern cannot be seen and few layers has also collapsed |  |
|  |  | 50 | The 3D printed structures demonstrate good printability, with the infill pattern distinctly visible and well-defined |  |
| 7 | 7 | 25 | Good printability but the infill pattern cannot be seen and few layers has also collapsed |  |
|  |  | 35 | The 3D printed structures demonstrate good printability, but the infill pattern cannot be seen and few layers has also collapsed |  |
|  |  | 50 | The 3D printed structures demonstrate good printability, with the infill pattern distinctly visible and well-defined layers was observed |  |

### 3.3 Analytical Characterization

The following components, namely SF, Gelatin Type A, GMA, MTGase, and SilkMA-Gelatin, were evaluated in FTIR and represented in **Figure 4**. Gelatin Type A exhibits broad peaks around ∼1729-1700 cm^-1^ (amide I/II band), 1729-1740 cm^-1^ (C=O stretching), ∼ 1530-1540 cm^-1^ (N-H stretch), ∼ 1230-1240 cm^-1^ (amide III), and ∼1050-1100 cm^-1^ (C-O stretching)^39^. MTGase showed similar amide peaks near ∼3280-3300 cm^-1^, ∼ 1650 cm^-1^, and ∼1530 cm^-1^ ^40^. The spectrum of GMA is marked by peaks around ∼1715-1725 cm^-1^ (C=O stretching) (green bands), ∼1635 -1640 cm^-1^ (C=C stretching), ∼950-1100 cm^-1^ (C-O-C stretching), and a distinct peak near ∼815-830 cm^-1^ associated with epoxy ring vibrations. SF displays the typical protein backbone peaks at ∼ 1650 cm^-1^(amide I), ∼1540 cm^-1^ (amide II), ∼1230 cm^-1^ (amide III), and ∼1070 cm^-1^ (C-O stretching)^41^. Upon Methacrylation, SilkMA retains the amide I, II, and III peaks and introduces new peaks at ∼1720 cm-1 (C=O stretching) and1635 cm^-1^ (C=C stretching)^42^. The SilkMA–gelatin composite shows combined features of both gelatin and SilkMA constituents, showing broad N-H/O-H stretching around ∼3300cm^-1^, strong amide I and II peaks (∼1650 and ∼1540 cm⁻¹) ^43^, the ester peak at ∼1720 cm⁻¹, the C=C signal at ∼1635 cm⁻¹, C–O stretching near ∼1040 cm⁻¹ and 2900–3000 cm⁻¹ (C-H stretch) blue band correlating it to change in methacrylation bonds. Orange band highlights the area around 1000-1200 cm^-1^ associated with C-O stretching and amide III groups which is corresponding with the SilkMA-Gelatin ink that contains MTGase. These spectral features confirm the successful incorporation and functionalization of each component.

**Figure 4:**
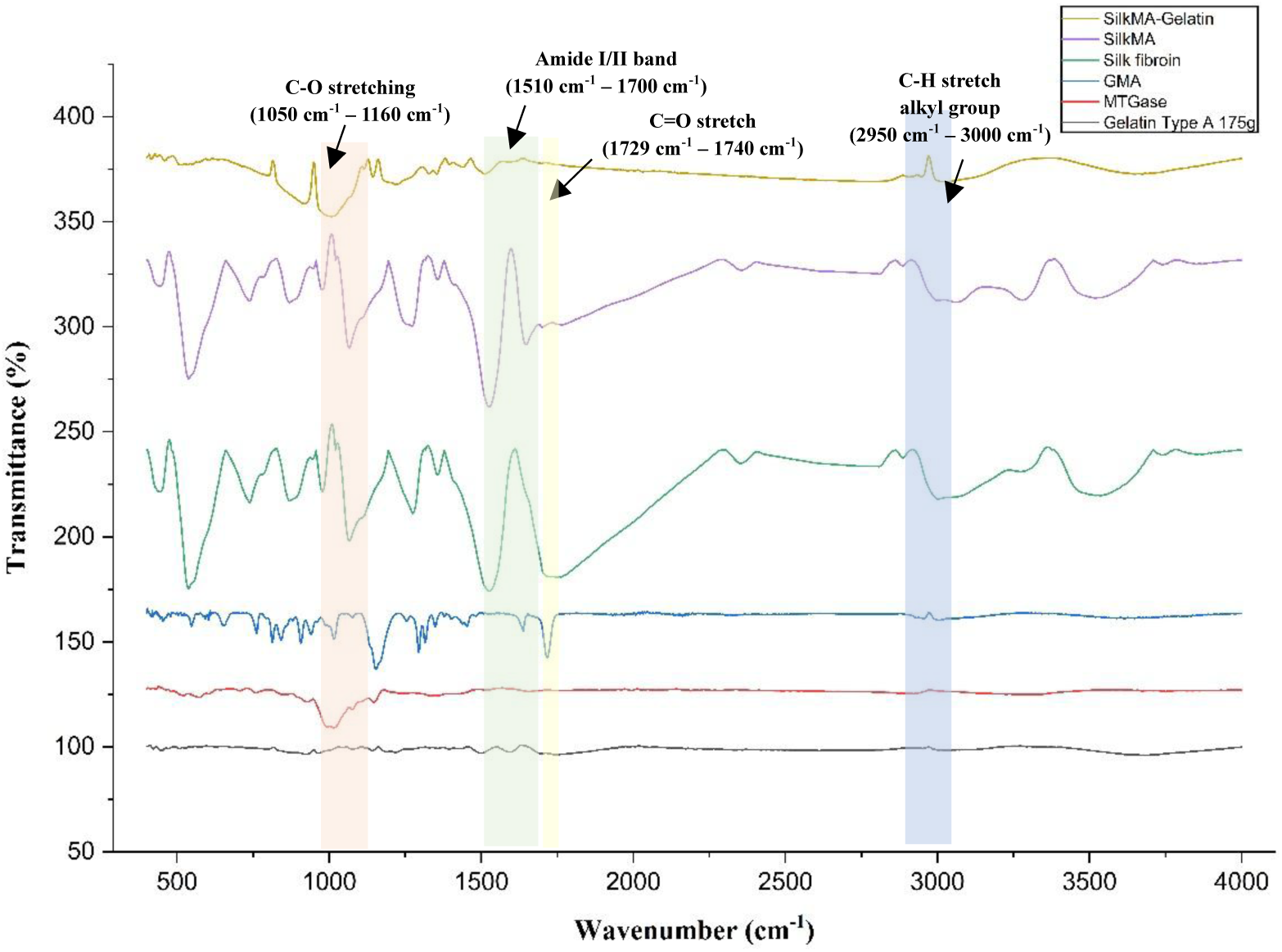
FTIR spectra of SF, SilkMA, SilkMA-Gelatin, GMA, MTGase and gelatin Type A 175G, Samples were analysed in the range of 500-4000 cm^-1^

SilkMA-Gelatin hybrid scaffolds contain overlapping amide bands due to the proteinaceous nature of both natural polymers, depicting a band near 3100-3700 cm⁻¹ (N-H and O-H band) indicating strong hydrogen bonding interactions between SilkMA and gelatin chains. According to Yung et al. peaks around 2850-2950 cm⁻¹ correspond to aliphatic C-H band which became more prominent after MTGase crosslinking which can be due to molecular rearrangement^44^. Similar peaks were observed around ∼2930-3000 cm^-1^ corresponding to presence of MTGase in SilkMA-Gelatin scaffold.

Shifting of amide I and II bands in silkMA as compared to silk fibroin indicates successful crosslinking by GMA thereby indicating successful crosslinking.

### 3.4 Thermal Characterization (DSC)

The DSC thermograms of gelatin, SilkMA, and SilkMA-Gelatin are provided. **Figure 5** presents important insights into their thermal transitions and structural stability over the temperature range of 30–350°C. Pure gelatin exhibited a glass-transition temperature (T_g_) near 50°C with a subsequent transition around 82°C corresponding to the melting of triple-helix domains **(Figure 5a)**. An intense endothermic peak at 168°C can be attributed to the gelatin denaturation, followed by a broad endothermic peak above 260°C, indicating decomposition, and an exothermic event near 330°C due to chain disintegration^45^. In contrast, SilkMA displayed a broad endothermic region between 44–150°C, attributed to water evaporation and conformational changes in water-plasticized chains **(Figure 5b)**. At elevated temperatures ∼203°C, it exhibits a broad glass-transition temperature followed by an endothermic peak at ∼289 °C and 300°C, which were associated with its decomposition and thermal degradation^46^. The SilkMA–gelatin blend **(Figure 5c)** exhibited a broad endothermic peak centered at 122 °C that reflected overlapping processes of water evaporation of silk, as well as melting, and partial denaturation of gelatin, followed by a glass transition at ∼192°C, which appeared at an intermediate between its components. At higher temperatures, a successive endothermic hump at 255°C and a peak at 271°C corresponded to the onset of SilkMA and gelatin degradation, respectively. Notably, the absence of the exothermic peak observed for pure gelatin at 332°C highlights the improved thermal stability of the SilkMA–gelatin blend. Overall, the DSC analysis shows that blending SilkMA with gelatin shifts the Tg to an intermediate value and suppresses the exothermic degradation peak of pure gelatin, thereby enhancing the thermal stability and structural integrity of the hybrid system. This improvement highlights the synergistic interaction between SilkMA and gelatin, making the blend a promising thermally robust biomaterial^47^.

**Figure 5:**
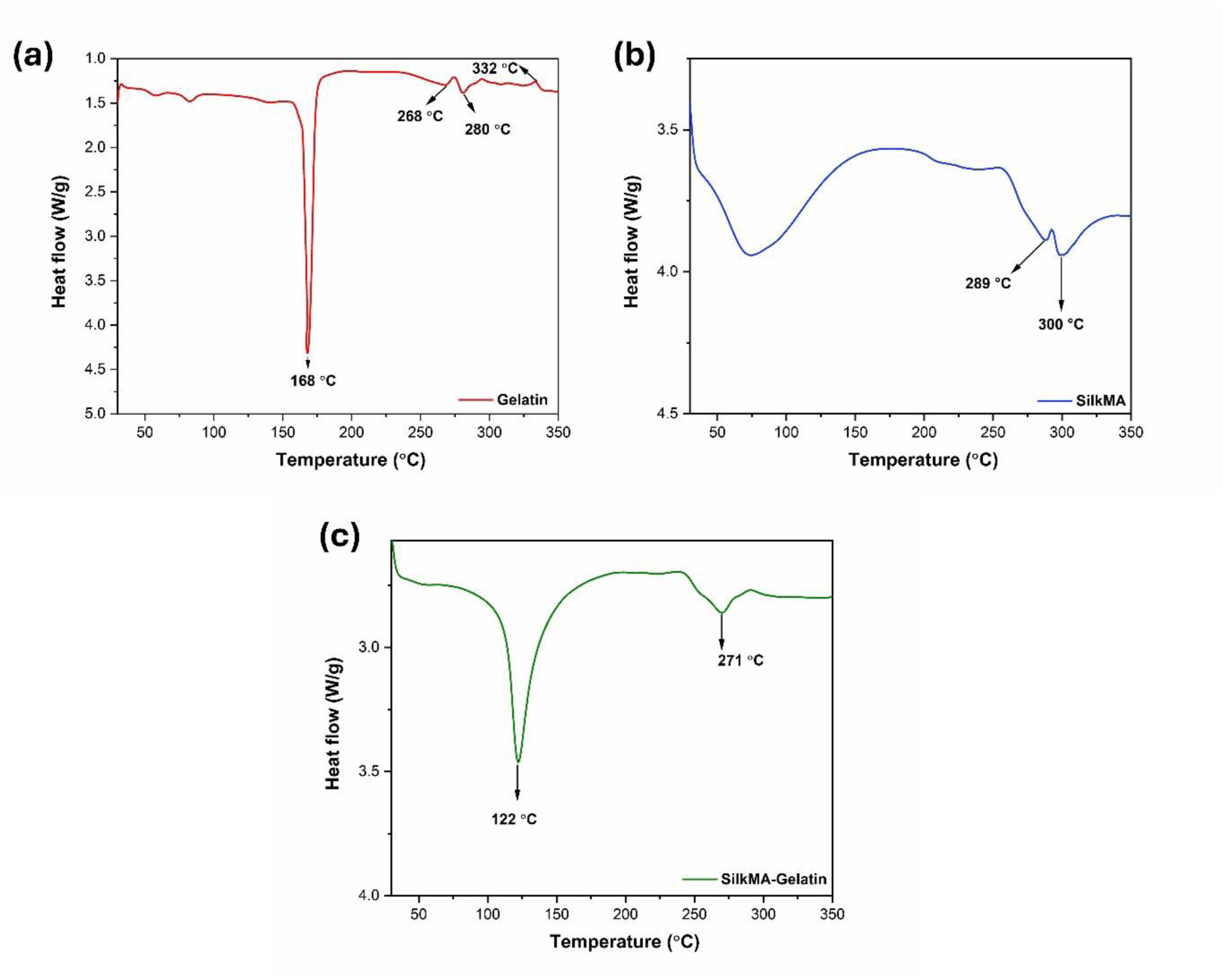
Differential scanning calorimetry (DSC) thermograms of (a) Gelatin, (b) SilkMA, and (c) SilkMA–Gelatin blend recorded over the temperature range of 50–350°C

### 3.5 Printing & CAD modelling

The 3D scaffolds were printed successfully, further demonstrating the functional versatility of the ink. The ink effectively created the uniform grid patterning, pores were interconnected, and consistent thickness throughout the 5 layers, conforming good interlayer bonding and structural uniformity **(Figure 6)**.

**Figure 6:**
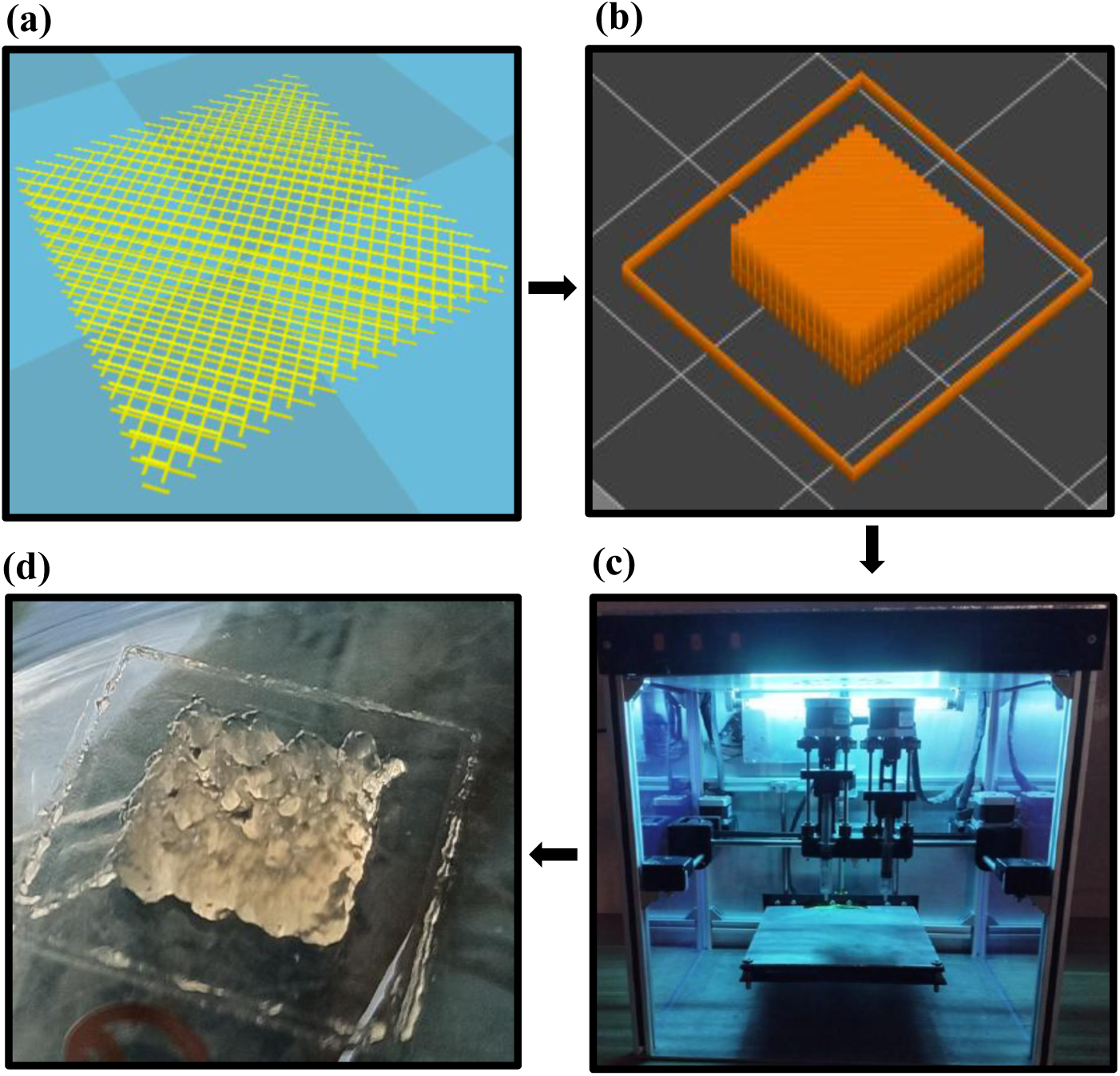
(a) 3D scaffold model was designed using Autodesk Fusion 360 for creating .stl file, (b) Sliced 3D model using Cura software for the generation of G-code, (c) 3D printing of the scaffold using Alfatek extrusion-based 3D bioprinter, (d) 3D printed scaffold

### 3.6 Rheological assessment of 3D SilkMA-Gelatin scaffold

The rheological properties of Silk–Gelatin and SilkMA–Gelatin ink were systematically investigated to evaluate their flow behavior and mechanical stability. The viscosity measurement performed over a shear rate range of 0.01 to 100s^-1^ revealed that both Silk-gelatin and SilkMA-Gelatin exhibited shear thinning behavior, where viscosity decreases with increasing shear rate, confirming their non-Newtonian nature **(Figure 7a)**. At low shear rates, Silk–Gelatin showed higher viscosity than SilkMA–Gelatin, indicating the presence of a denser physical network. However, at higher shear rates, the viscosities of both systems converged due to structural breakdown and polymer chain alignment under shear. Notably, at a shear rate of 50s^-1^, the apparent viscosity of Silk-Gelatin is 37,363 mPa.s, which is comparatively higher than SilkMA-Gelatin, which is 23,434 mPa.s, indicating easier flow through the nozzle and reduced extrusion pressure during bioprinting. This lower viscosity while maintaining structural cohesion suggests that SilkMA-Gelatin is better suited for gentle cell encapsulation and high-resolution printing^48^.

**Figure 7:**
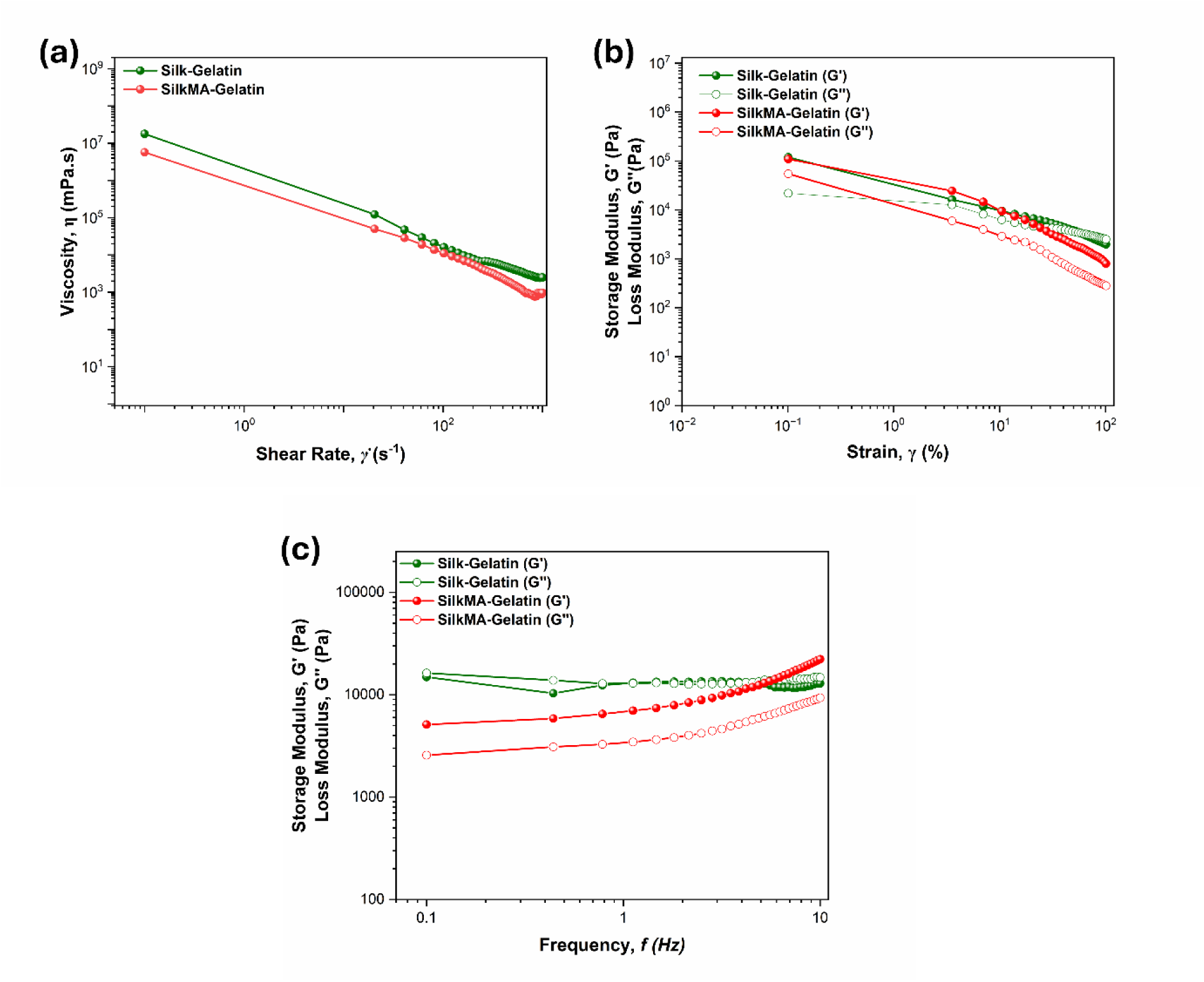
(a) Shear-dependent viscosity of Silk-Gelatin and SilkMA-Gelatin, (b) oscillatory amplitude sweep at a constant frequency of oscillation of 1Hz, and (c) frequency sweep in the linear viscoelastic range (0.1-10 Hz) at a constant strain of 0.5%

The oscillatory strain sweep measurements **(Figure 7b)** further demonstrated that both materials possessed a well-defined linear viscoelastic region (LVR) at low strain, where the storage modulus (G′) exceeded the loss modulus (G″), reflecting dominant elastic, gel-like behavior^49^. With increasing strain, both G′ and G″ decreased, indicating progressive network disruption and eventual structural failure at high deformation. Further, the oscillatory frequency sweep tests performed over 0.1 to 10 Hz at constant strain of 0.5% **(Figure 7c)** show that SilkMA–Gelatin maintained G′ consistently higher than G″ across the frequency range, reflecting stable gel-like behavior with improved elasticity and structural integrity compared to Silk–Gelatin, thus suitable for 3D bioprinting^50^.

### 3.7 Characterization of SilkMA-Gelatin scaffolds

#### 3.7.1 Microscopic analysis of SilkMA-Gelatin scaffold

The microstructure of silkMA scaffolds was observed by SEM. The images revealed interconnected porous microstructures throughout the scaffold to favour cell infiltration and proliferation ^51^ **(Figure 8a)**. It was observed that the pore size of the 3D printed scaffold ranged from 5.07 μm to 11.6 μm ^52^**(Figure 8b)**, highlighting the wide distribution of the pores throughout the scaffolds. Such porosity is critical for cell infiltration, nutrient transport and waste diffusion which collectively enhances tissue integration and regeneration^53^. Smaller pore sizes in this range are known to facilitate fibroblast attachment and proliferation as well as to promote ECM deposition during early stages of wound healing^54^. The interconnected morphology achieved through the optimised SilkMA-Gelatin formulation and 3D printing parameters ensures homogeneous cell migration and vascular infiltration within the construct^54^.

**Figure 8:**
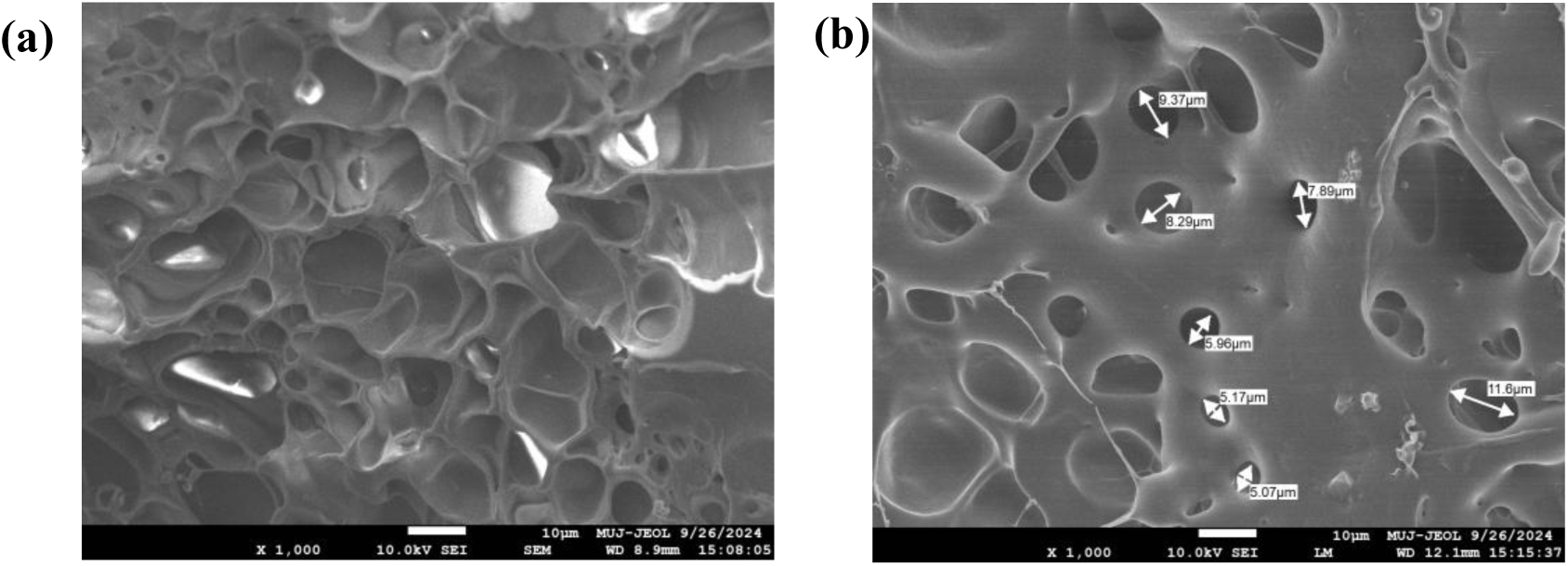
Microscopic analysis of the internal architecture of 3D printed SilkMA–Gelatin scaffolds using FE-SEM (a) Image showing a highly interconnected porous structure, and (b) image depicting pore size distribution at the same magnification

#### 3.7.2 Swelling property

As shown in **Figure 9**, PBS and the DMEM environment demonstrated swelling of the 3D printed scaffold, with significant swelling observed in PBS compared to DMEM at all time points. Within the first 30 minutes, 3D printed scaffolds in PBS showed a sharp increase in swelling ratio, reaching nearly 80 ± 1.1 % whereas scaffolds in DMEM exhibited a gradual rise of about 45 ± 3.5 %. Over the next few hours, swelling in PBS continued to increase, peaking around 89.4 ± 0.5 % after 24 hours, after which a slight decrease was observed. On the contrary, scaffolds immersed in media exhibited a steady, gradual increase, stabilizing around a 64 ± 2.1 % swelling ratio by 96 hours.

**Figure 9:**
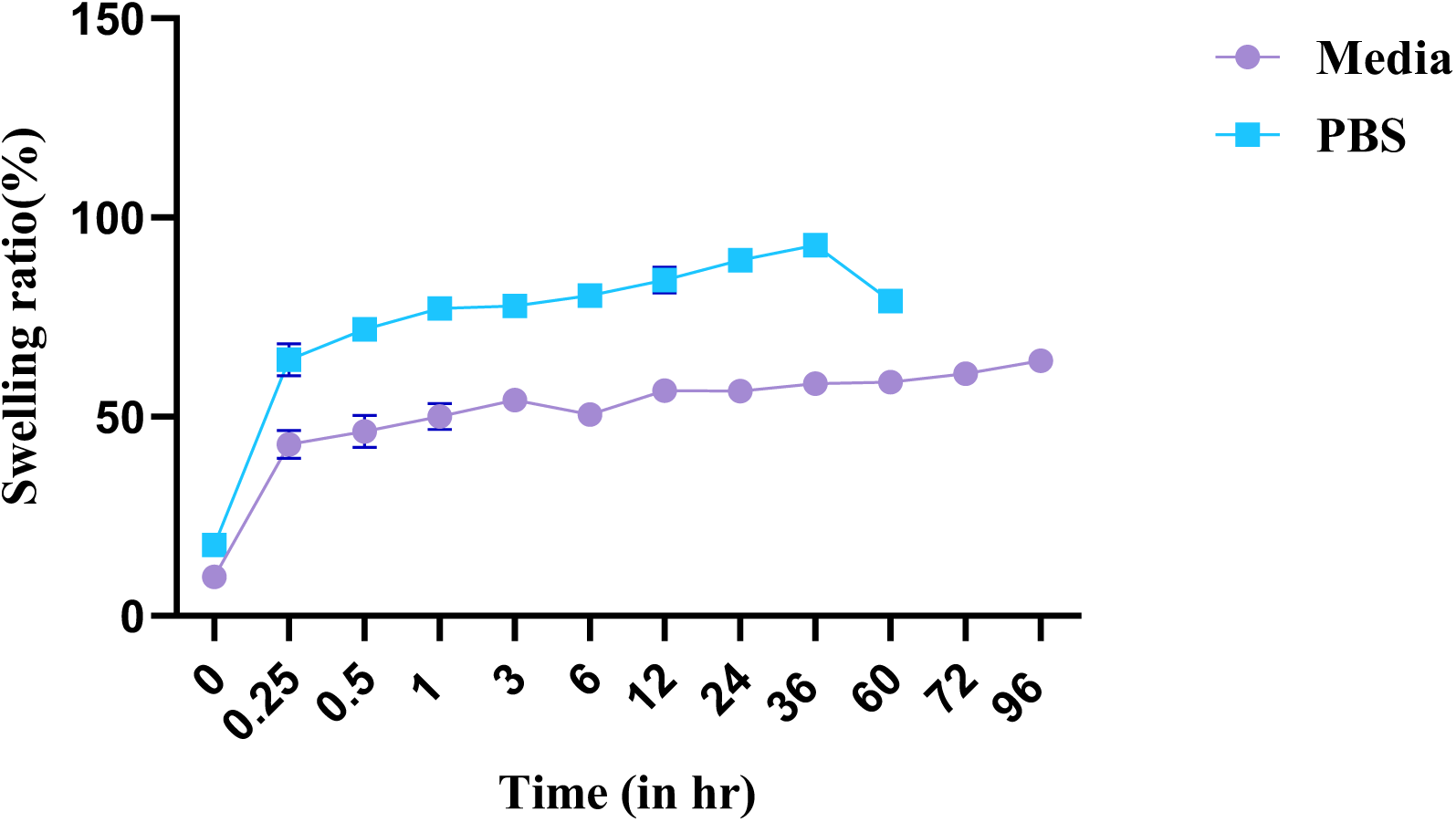
Swelling behavior of 3D printed SilkMA-Gelatin scaffold in PBS (pH-7.4) & media (DMEM) at 37°C as a function of time

The higher swelling in PBS can be attributed to osmotic pressure differences and the absence of serum proteins which allow restrained diffusion of water molecules into the polymeric matrix, whereas DMEM contains amino acids, glucose and proteins that interact with the polymer network through hydrogen bonding or ionic screening effectively reducing the free volume and restricting excessive fluid uptake^55^.

#### 3.7.3 Wettability analysis

The contact angle provides critical insights on the hydrophilicity of the 3D printed scaffolds for assessing the materials’ suitability, which decides cell adhesion. It was observed that the SilkMA-Gelatin scaffold exhibits remarkable hydrophilic properties, ranging from 69.5° ± 4.5 to 82° ± 1.4, as shown in **Figure 10**. Furthermore, employing the Drop Analyzer Software, it was assessed that the droplet volume remaining on the surface of the 3D printed scaffold was less than the targeted 50μL. This intriguing observation highlights the impressive ability to absorb liquid, reinforcing its potential as a promising material for biomedical applications that demand effective cell interactions^56^.

**Figure 10:**
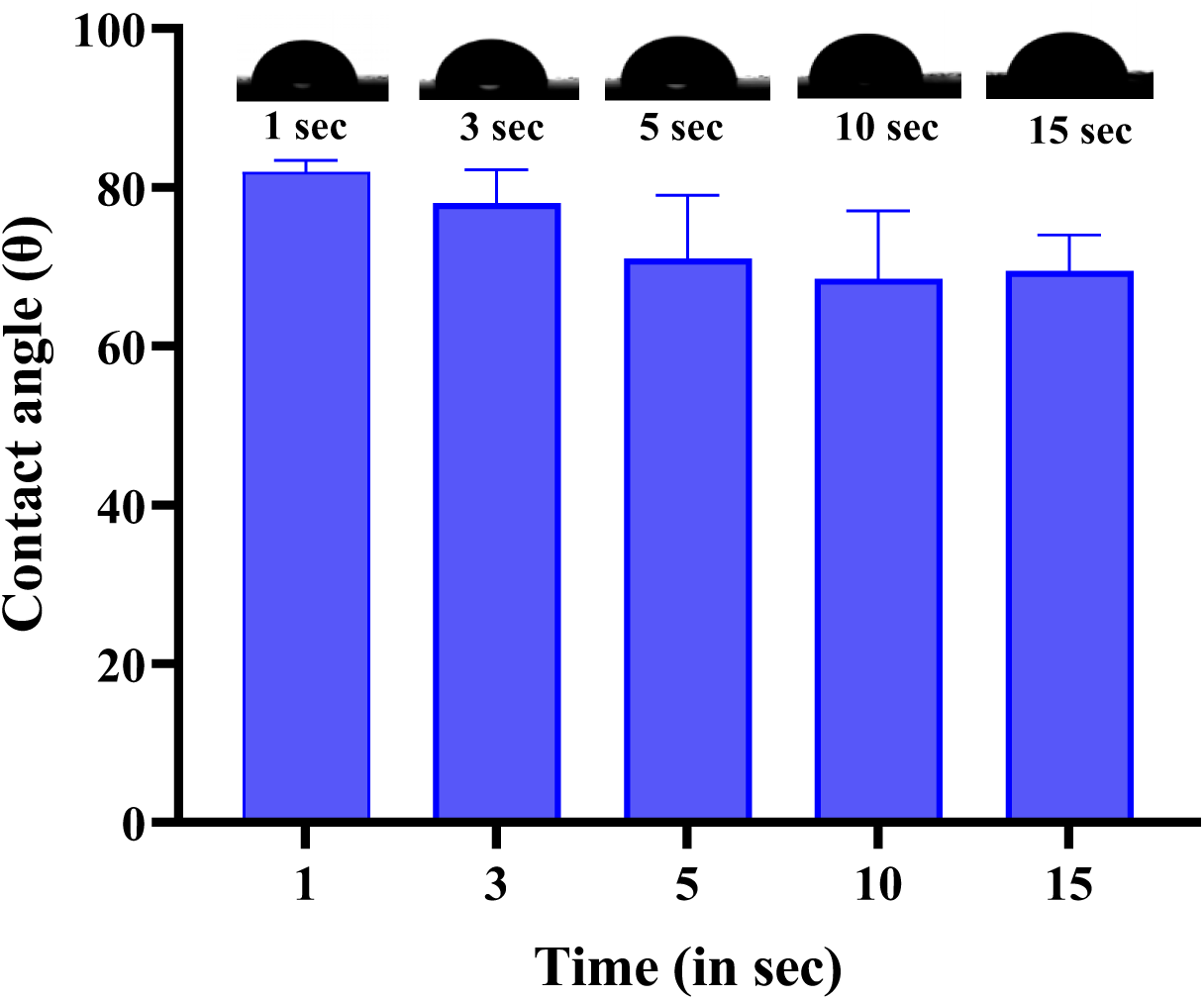
Contact angle of 3D printed scaffold. Representative image of the water droplet over the scaffold surface

### 3.8 Cytocompatibility assays of SilkMA-Gelatin *scaffolds*

#### 3.8.1 Live/dead assays

Live/dead staining images revealed the viability and distribution of NIH3T3 fibroblasts cells encapsulated within the 3D bioprinted SilkMA-Gelatin scaffolds over time. On Day 1, high density of live cells (green) was observed, indicating improved cell viability and successful encapsulation post-printing, as shown in **Figure 11a**. By Day 7, cells appear more distributed, depicting enhanced cell attachment and proliferation within the 3D scaffold **(Figure 11b)**. By Day 14, the majority of cells remain viable, indicating sustained cytocompatibility of the 3D scaffold in **Figure 11c**. The overall trend suggested that the 3D bioprinted scaffold supports cell survival and attachment over two weeks.

**Figure 11:**
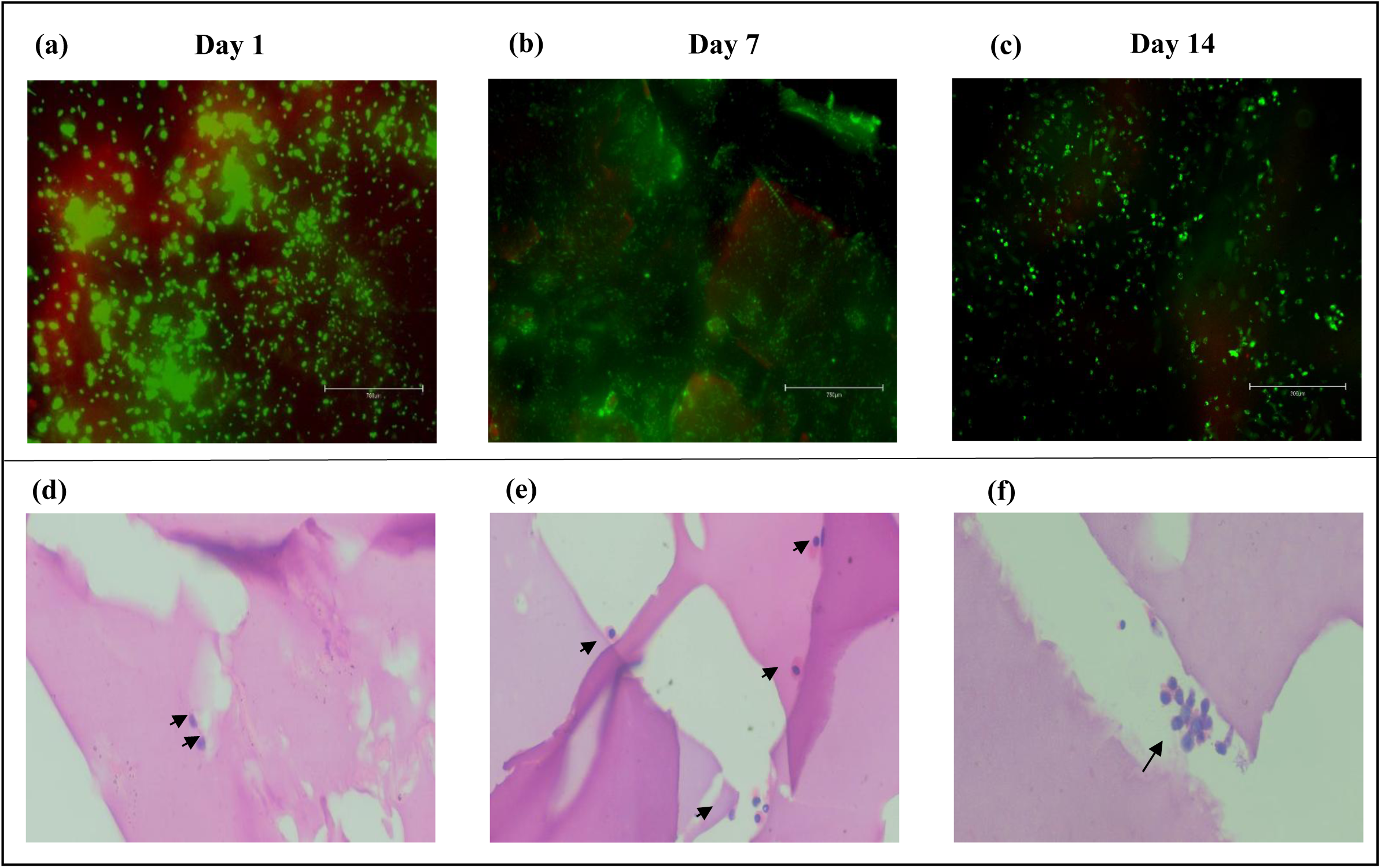
Cytotoxicity of SilkMA-Gelatin solutions: Live/Dead images of 3D scaffolds bioprinted using NIH3T3 on (a) Day 1, (b) Day 7 and (c) Day 14; Histological characterization of 3D bioprinted scaffolds on (d) Day 1, (e) Day 7 and (f) day 14 at 40X magnification

#### 3.8.2 H&E staining

H&E staining reveals the cell distribution and matrix interaction within the 3D bioprinted SilkMA-Gelatin scaffolds over time, as shown in **Figure 11d**. On Day 1, minimal number of cells are observed (black arrows), mainly localized around the scaffold edges, indicating initial cell adhesion. By Day 7, multiple cells are evident, with some appearing infiltrated within the scaffold pores, suggesting cell migration and improved interaction with the scaffold matrix [69]. On Day 14, the number of embedded cells increases further, and cells are more evenly distributed across the scaffold, indicating enhanced biocompatibility and cell proliferation over time. The progressive increase in cellular presence proves that SilkMA-Gelatin as a supportive material for 3D bioprinting applications.

### 3.9 Biocompatibility assays of SilkMA-Gelatin scaffolds

An increase in body weight was observed in Wistar rats at all the time points post-subcutaneous implantation of the 3D scaffold, as shown in **Figure 12a**, suggesting the absence of post-implantation stress, malnutrition, or severe complications. Mean body weights of SD rats implanted with 3D printed SilkMA-Gelatin slightly increased from 14 days and 1 month. There was no statistically significant difference between the groups, which indicates favorable host response and absence of chronic inflammation or immune rejection ^57^.

**Figure 12:**
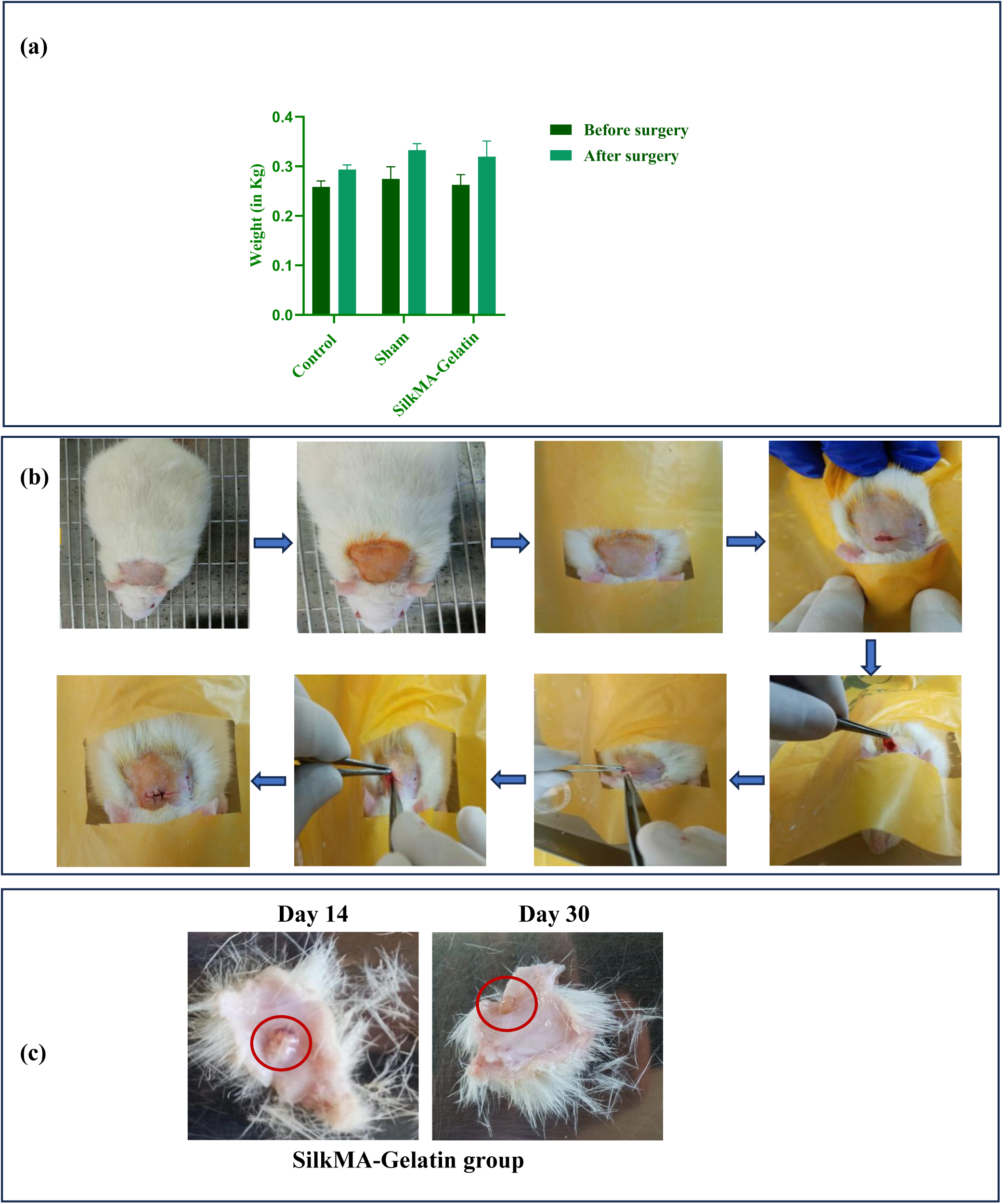
(a) Body weight of Wistar rats before (9 weeks) and after (11 weeks & 13 weeks) surgery, (b) Implantation process of putting the 3D printed SilkMA-Gelatin scaffold inserted inside the subcutaneous pocket, (c) 3D printed scaffolds post-implantation

Implanted scaffolds retrieved after 14 days and 1 month demonstrated in **(Figure 12c)**, have shown structural persistence and tissue integration, highlighting the long-term stability and biocompatibility of the constructs. The partial disintegration of the 3D printed SilkMA-Gelatin scaffold signifies controlled degradation and successful interaction with the surrounding tissues. Presence of the scaffolds after day 14 and day 30 of the implantation suggests that the chemical and enzymatic crosslinking provided adequate mechanical stability to resist premature resorption while also allowing the gradual infiltration of the cells and remodelling of the skin^58^.

#### 3.9.1 Hematology

Hematological parameters were evaluated between day 0 (pre-treatment), the control (no treatment), and days 14 and 30 of post-implantation to assess systemic biocompatibility and potential inflammatory response induced by the implanted scaffolds. In **Figure 13a**, WBC count of SilkMA-Gelatin (13.15 ± 0.21 × 10³/µL) and sham (11.55 ± 2.19 × 10³/µL) is slightly higher as compared to control group (8.35 ± 0.35 × 10³/µL). Around Day 30 the values of the treatment group (SilkMA-gelatin) come within the normal physiological range of 8.40 x 10^3^/μL with no statistically significant difference observed.

**Figure 13:**
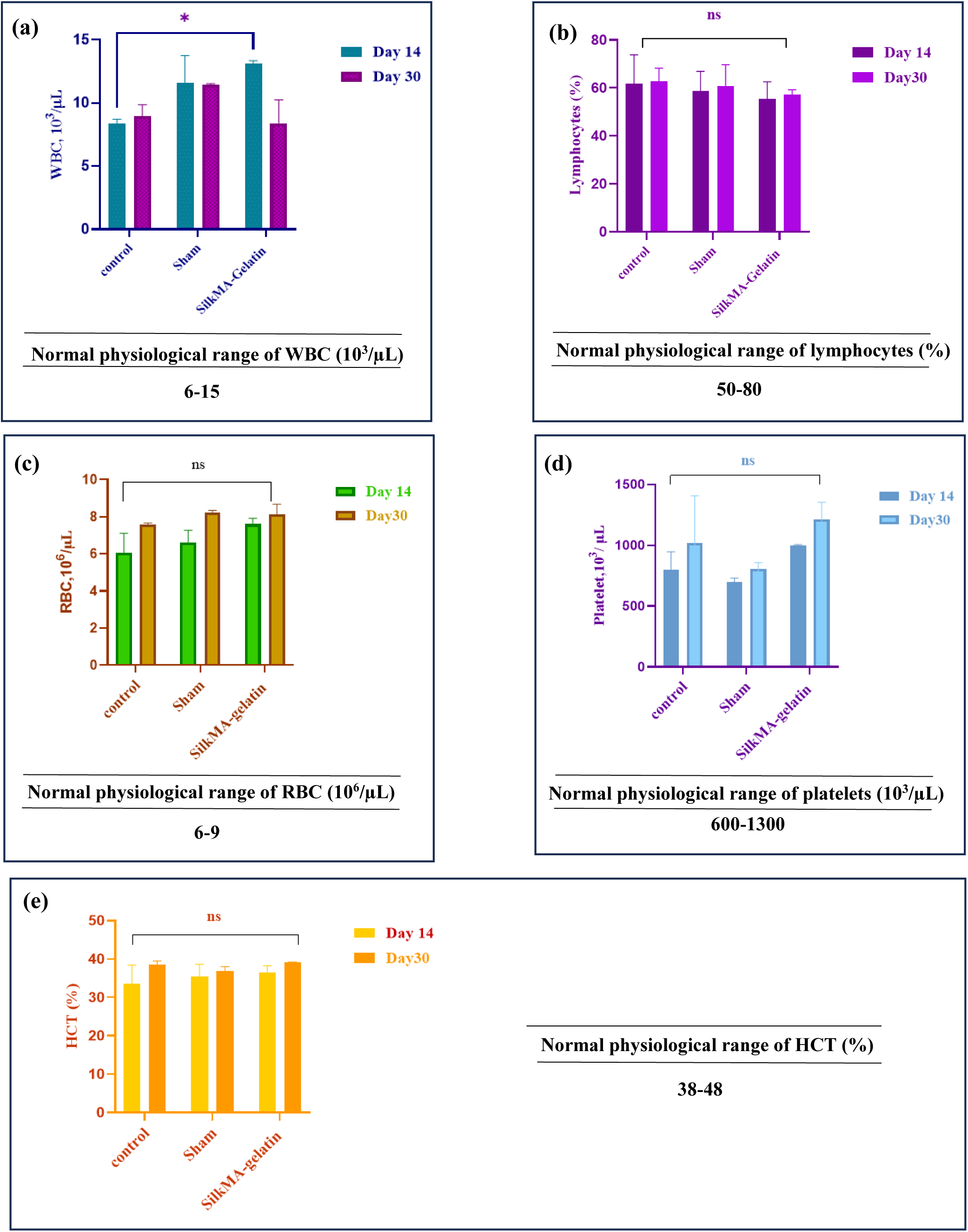
Complete blood count data after day 14 and day 30 of implantation for control, sham and SilkMA-Gelatin group. CBC parameters: (a) WBC count (10^3^/µL), (b) Lymphocytes (%), (c) RBC count (10^6^/µL), (d) Platelet count (10^3^/µL), (e) HCT (%)

**Figure13b**, illustrates lymphocytes percentage on day 14 and day 30 across groups ranging between 55.5 ± 7.07% (SilkMA-Gelatin), 58.70 ± 8.20% (Sham) and 61.80 ±12.02% (control) for day 14 and no appreciable change was observed at day 30 (variation of 0.90 from day 14 across groups) suggesting no chronic immune activation.

In **Figure 13c**, at day 14 the RBCs count of SilkMA-Gelatin (7.60 ± 0.31 x 10^6^/µL), sham (6.63 ± 0.64 x 10^6^/µL) and Control (6.07 ± 1.04 x 10^6^/µL) which is within the range of normal count. On day 30, the RBCs count has increased by a variation of 8.0 ± 0.4 x 10^6^/µL that is also within the normal range suggesting no abnormality related to RBC count.

In **Figure 13d**, Platelet count of SilkMA-Gelatin group (1000 ± 5.65 x 10^6^/µL showed an increase on day14 as compared to the sham (701.00 ± 29.69 x 10^6^/µL) and control group (799.5 ± 146.3 x 10^6^/µL but this increase is still within the normal range. At day 30, the trend changed and the counts came at a comparable range between control (1018 ± 387.49 x 10^6^/µL) and the treatment group (1212.5 ± 140.714 x 10^6^/µL) whereas count for sham was still around 804 x 10^6^/µL. However, all values remained within the normal range.

In **Figure 13e**, HCT percentage is shown for day 14 and 30. It was observed that there were no significant changes within the groups. For day 14, SilkMA-Gelatin (36.45 ± 1.76 x 10^6^/µL), Sham (35.50 ± 3.11 x 10^6^/µL) and control (33.65 ± 4.73 x 10^6^/µL) remain within the normal range. Similarly, for day 30 the values have slightly increased for all the groups, i.e, 39.15 ± 0.070 x 10^6^/µL for treatment, 36.80 ± 1.27 x 10^6^/µL for sham and 38.50 ± 0.98 x 10^6^/µL for control; however, the variation is non-significant.

#### 3.9.2 Serum biochemistry

##### 3.9.2.1 Alkaline phosphatase (ALP) assay

The enzymatic activity levels, measured in µmol/min/mL, were evaluated in control, sham, and SilkMA-Gelatin implanted groups at days 14 and 30. As shown in **Figure 14a**, on day 14, the control group showed an average activity of 3 ± 0.89 µmol/min/mL, the sham group 2.9 ± 0.72 µmol/min/mL, and the SilkMA-Gelatin group 2.32 ± 0.042 µmol/min/mL. By day 30, all groups showed a slight reduction in activity, with control and groups measured around 2 ± 1.19 µmol/min/mL and 2.5 ± 0.13 µmol/min/mL, respectively, while the SilkMA-Gelatin group remained consistent at 2.4 ± 0.06 µmol/min/mL. No notable differences were observed between them. Statistical analysis using two-way ANOVA revealed no significant differences among the groups or between time points, indicating that the 3D printed scaffold did not induce adverse enzymatic response over time and maintained the normal ranges of blood ALP levels along with the control and sham groups^59^.

**Figure 14.**
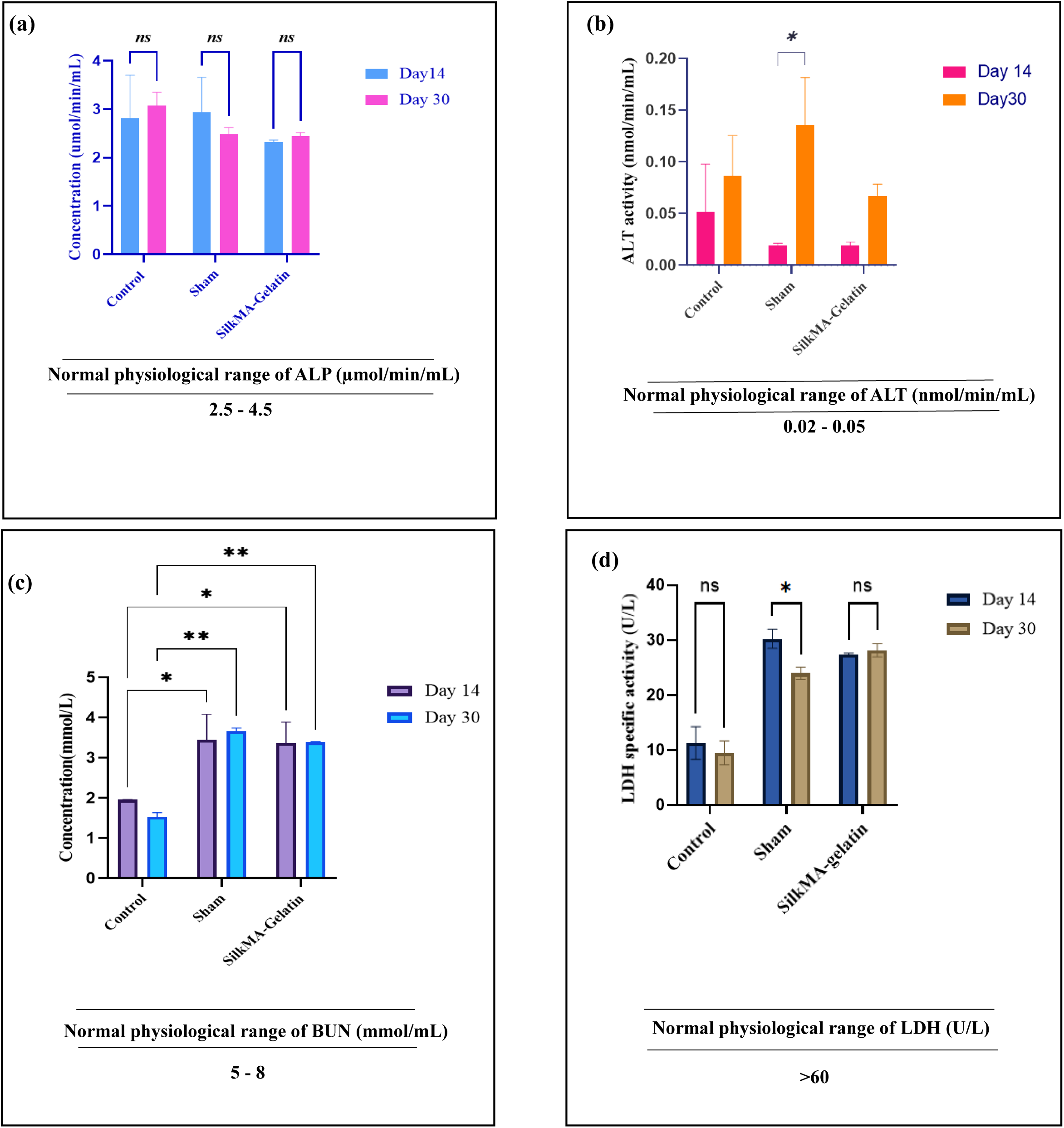
(a-d): *In vivo* biocompatibility of SilkMA-Gelatin scaffolds. Changes in (a) ALP, (b) ALT, (c) BUN and (d) LDH suggest that the 3D printed scaffolds had no hepatotoxicity and nephrotoxicity effects *in vivo.* For statistical analysis was performed using two-way ANOVA [*p≤0.05, **p≤0.01]

##### 3.9.2.2 Alanine Transaminase (ALT) assay

At day 14, all groups exhibited low ALT levels with no significant differences observed among the control, sham, and SilkMA-Gelatin groups. However, by day 30, ALT levels were observed to show a significant increase i.e.0.1353 ± 0.046 nmol/min/mL, as compared to what was detected in the sham group on day 14, i.e., 0.018953 ± 0.002052 nmol/min/mL, suggesting a potential hepatic response to surgical stress or local inflammation in the absence of the scaffold. SilkMA-Gelatin group showed no significant change in ALT levels between day 14 (0.018953 ± 0.00335 nmol/min/mL) and day 30 (0.066751 ± 0.011525 nmol/min/mL), indicating that the scaffold did not elicit a hepatotoxic response^59^. The control group maintains stable ALT levels throughout the study, ranging from 0.051646 ± 0.046235 nmol/min/mL to 0.086268 ± 0.039126 nmol/min/mL. These findings suggest that the 3D scaffold is biocompatible in terms of liver function and does not induce systemic hepatic toxicity ^60^, as shown in **Figure 14b**.

##### 3.9.2.3 Urea (BUN) assay

The urea concentration was measured across the control, sham, and SilkMA-Gelatin groups at day 14 and day 30, as shown in **Figure 14c**. On day 14, the control group showed the lowest urea level at approximately 1.96 ± 0.004 mmol/L, while the sham group exhibited urea levels of 3.45 ± 0.64 mmol/L. The SilkMA-Gelatin groups showed levels around 3.3 ± 0.52 mmol/L. By day 30, the urea level in the control group had decreased slightly to 1.5 ± 0.10 mmol/L, whereas the sham group showed a slight increase to 3.6 ± 0.07 mmol/L, and the SilkMA-Gelatin group remained consistent around 3.39 ± 0.08 mmol/L.

Statistical analysis using two-way ANOVA revealed no significant difference between the sham and SilkMA-Gelatin groups (p>0.05), indicating the SilkMA-Gelatin scaffold maintains urea levels comparable to physiological conditions.

##### 3.9.2.4 Lactate dehydrogenase (LDH) assay

As shown in **Figure 14d**, the control group exhibited low LDH activity at both time points, ranging from approximately 10-12 U/L, indicating minimal baseline cell damage, whereas the sham group showed a significant increase in LDH activity as compared to the control, at both 14 and 30 days, which is approximately 30-32 U/L, which can be due to surgical intervention.

LDH levels of the SilkMA-Gelatin group showed moderate levels of approximately 26-28 U/L, indicating reduced cytotoxicity compared to sham, reflecting good biocompatibility.

### 3.9 H&E Staining of Skin Post-Implantation

As shown in **Figure 15**, H & E staining of skin tissues post-implantation revealed progressive tissue remodeling across all groups. In the control group, the skin architecture remained intact with clearly defined epidermal and dermal layers along with visible hair follicles and sebaceous glands, indicating normal histology at both day 14 **(Figure 15a)** and day 30 **(Figure 15b)**. The Sham Group exhibited mild epidermal thickening and slight infiltration on day 14, which subsided by day 30, showing improved re-epithelialization and restoration of dermal structures.

**Figure 15:**
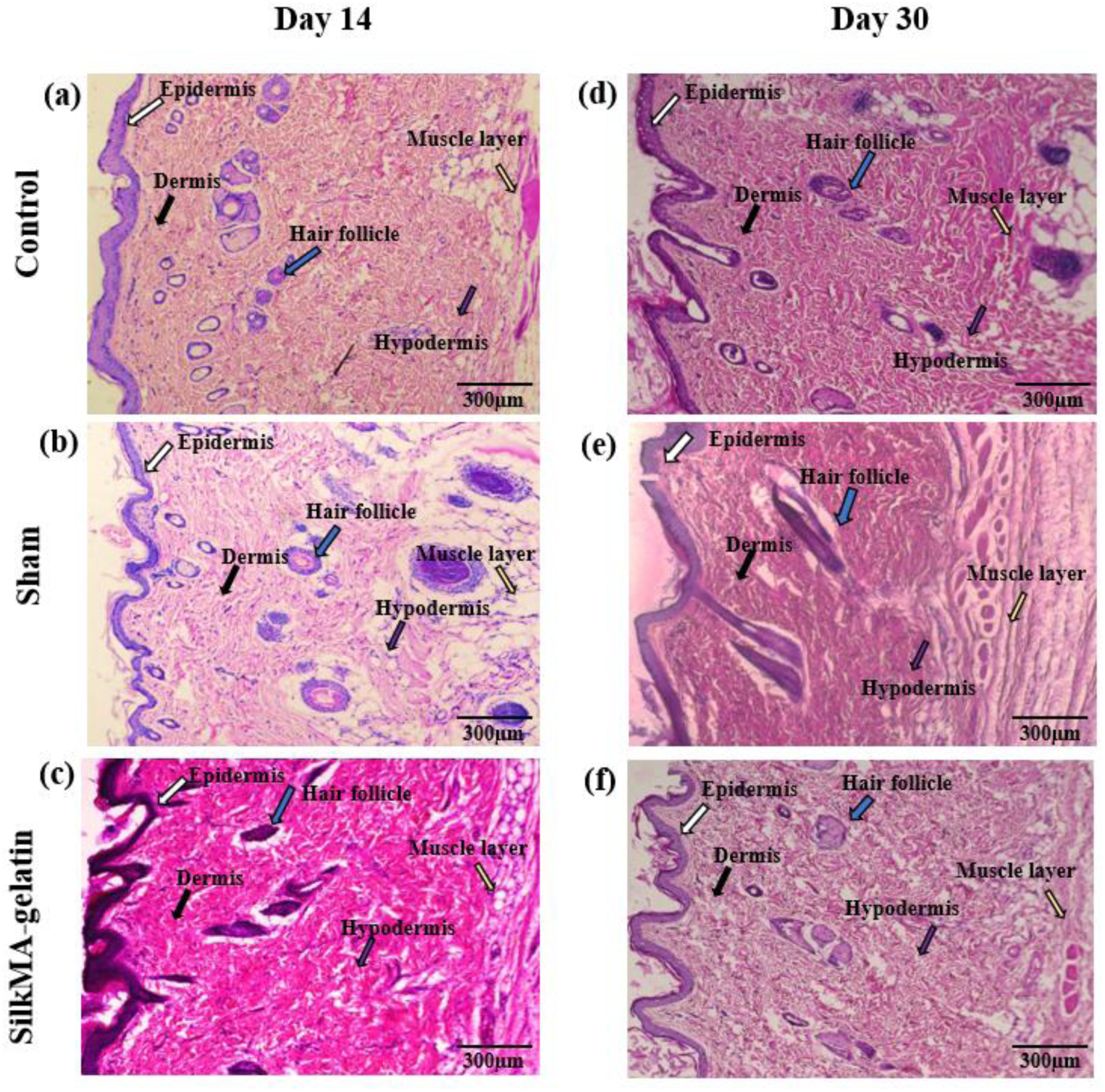
Biocompatibility assessment of scaffolds following subcutaneous implantation in Wistar rats, evaluated by H&E staining, Representative H&E-stained images of surrounding skin tissue at (a) control, (b) sham, and (c) 3D printed scaffold implantation sites on day 14, and (d) control, (e) sham, and (f) 3D printed scaffold implantation sites on day 30

The SilkMA-Gelatin group showed moderate inflammation and partial re-epithelialization on day 14. By the 30th day, the tissue exhibited better-organized epidermal layers, reduced inflammation, and the reappearance of skin appendages, such as hair follicles.

The short-term *in vivo* results validated that 3D printed scaffold poses no hindrance towards the regenerative potential of skin. **Table 3**, summarizes the histological evaluation of skin at day 14 and day 30 it was revealed that the regenerated skin where the 3D printed SilkMA-Gelatin scaffold was implanted displayed a well-defined epithelial layer, dermis and hypodermis along with the presence of skin appendages and minimal scar formation signifying angiogenesis and restoration of follicular units, which are key hallmarks of physiological healing. It was observed that after the implantation within 14^th^ day the hair growth of the rats began and there was minimal to no scar observed hinting towards functional skin regeneration.

**Table 3:**
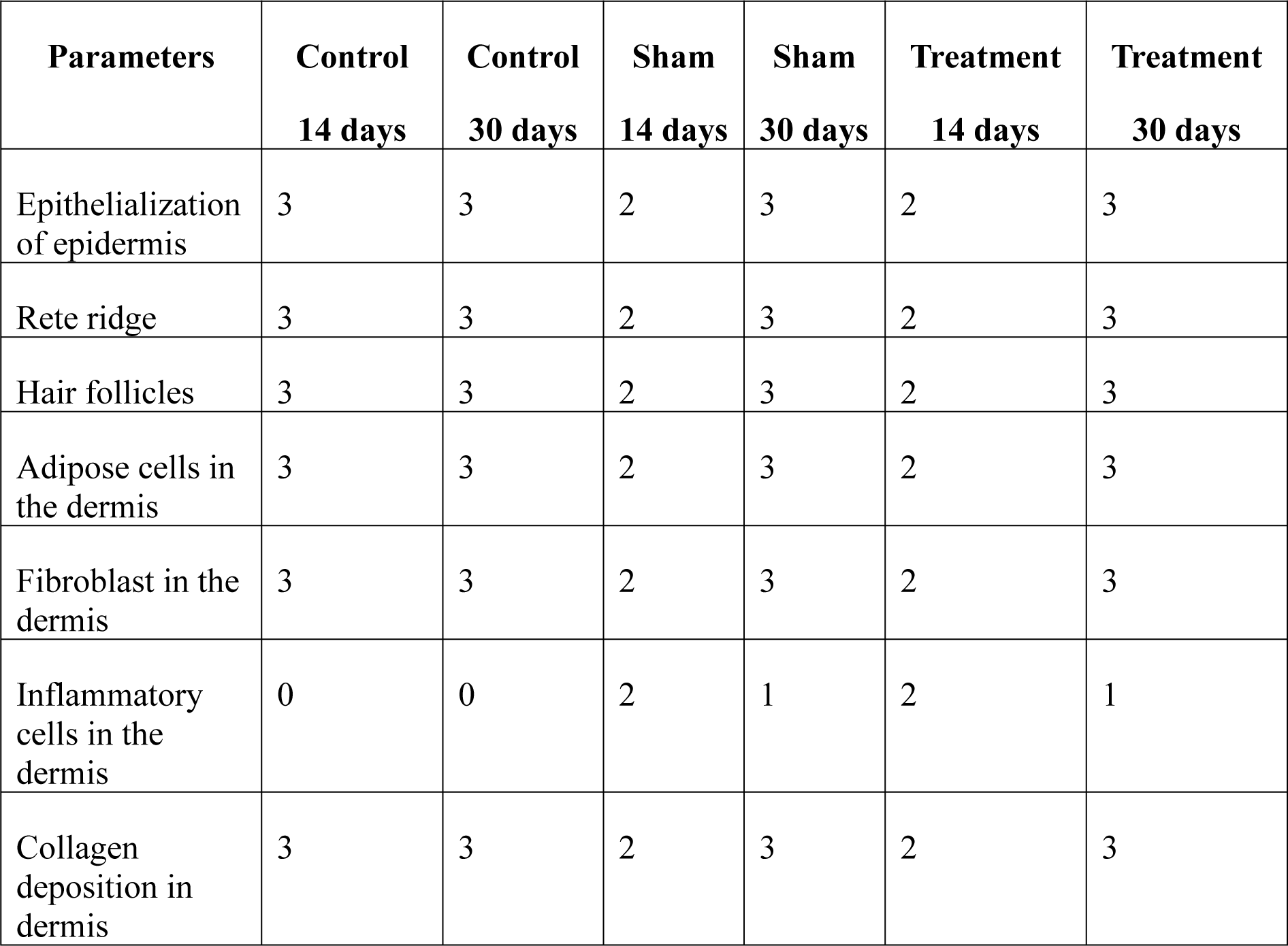
Scoring of rat skin on a scale of (0-3), 0 - absent; 1- mild; 2 - moderate; 3 – severe

| Parameters | Control<br>14 days | Control<br>30 days | Sham<br>14 days | Sham<br>30 days | Treatment<br>14 days | Treatment<br>30 days |
| --- | --- | --- | --- | --- | --- | --- |
| Epithelialization of epidermis | 3 | 3 | 2 | 3 | 2 | 3 |
| Rete ridge | 3 | 3 | 2 | 3 | 2 | 3 |
| Hair follicles | 3 | 3 | 2 | 3 | 2 | 3 |
| Adipose cells in the dermis | 3 | 3 | 2 | 3 | 2 | 3 |
| Fibroblast in the dermis | 3 | 3 | 2 | 3 | 2 | 3 |
| Inflammatory cells in the dermis | 0 | 0 | 2 | 1 | 2 | 1 |
| Collagen deposition in dermis | 3 | 3 | 2 | 3 | 2 | 3 |

### 3.11 H & E staining of the implanted 3D printed scaffold

As shown in **Figure 16**, H & E staining of the implanted 3D printed SilkMA-Gelatin scaffold revealed the presence of cellular infiltration and host tissue interaction with the biomaterial. The formation of fibrous tissue surrounding the 3D scaffold implies an early stage of tissue remodeling. Cellular presence within the 3D printed SilkMA-Gelatin scaffold provides evidence of its biodegradability and capability of supporting cell migration, making it a promising candidate for 3D bioprinting using an extrusion-based 3D bioprinter.

**Figure 16:**
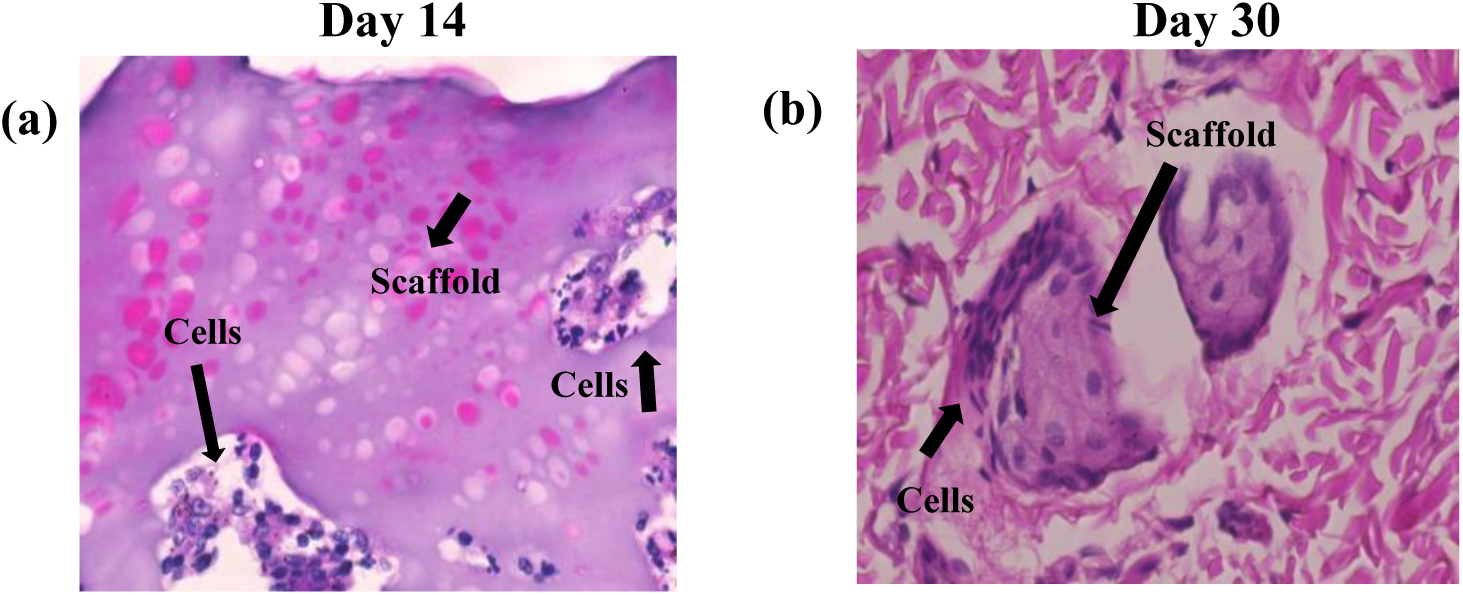
Histological evaluation of 3D printed SilkMA-Gelatin scaffold integration and cellular infiltration at 60x magnification on (a) day 14 and (b) day 30 post-implantation using H&E staining

Infiltration of the cells within the scaffolds at day 14^th^ and day 30^th^ is an indication of the hospitable environment around the implanted area which helped the cells to infiltrate, proliferate and regenerate the implanted area. This biocompatibility reflects that the scaffold provides optimized porosity and nutrient diffusion for the cells to proliferate^61^.

## Conclusion

A group of components were processed together to develop an innovative silk-based ink, namely SilkMA-Gelatin highlighting the functional and biocompatibility. Establishing SilkMA-Gelatin as a promising biomaterial for extrusion 3D printing. The integration of GMA, Gelatin and MTGase resulted in a structurally stable, porous, cytocompatible as well as biocompatible 3D printed scaffold. Presence of increasing number of live cells from day 1 to day 14 in live/dead assay and H&E staining established the cytocompatibility of the 3D printed scaffold. Also, healing of the area of implantation without any scarring and absence of any toxicity in *in vivo* studies establishes biocompatibility of the 3D printed SilkMA-Gelatin scaffold. The developed ink can be used along with various cytokines and cells to develop a 3D printed/bioprinted wound healing scaffold to address various types of wounds. The extrusion 3D bioprinting will have a profound effect to allow maximum cell survival post-printing and will have promising effects in regenerative medicine.

## Author contribution

**Kirthanashri S. V.**: conceptualization, investigation, data curation, writing– original draft, editing, reviewing and validation; **Prachi Agarwal**: investigation, data curation, visualization, writing– original draft; **Jagnoor Singh Sandhu:** Surgical help during implantation process; **Himanshu Kumar Bhatt:** data analysis for analytical & thermal characterization **Paramita Das:** data analysis for analytical & thermal characterization; **Raviraja N Seetharam:** editing, reviewing and validation

## Conflicts of interest

The authors declare no competing interests.

## Acknowledgment

The authors would like to thank the Department of Biotherapeutics Research (DBR), MAHE, Manipal for research and infrastructural support.

## Notes

### Competing Interest Statement

The authors have declared no competing interest.

